# The Rice Plastidial Phosphorylase Participates Directly In Both Sink And Source Processes

**DOI:** 10.1101/2020.07.09.191585

**Authors:** Kaan Koper, Seon-Kap Hwang, Magnus Wood, Salvinder Singh, Asaph Cousins, Helmut Kirchhoff, Thomas W. Okita

## Abstract

A distinctive structural feature of the higher plant plastidial starch phosphorylase (Pho1) is a 50 to 82 amino acid long peptide (L50 - L82), which is absent in phosphorylases from non-plant organisms. To study the function of the rice Pho1 L80 peptide, we complemented a *pho1^−^* rice mutant (BMF136) with the wildtype Pho1 gene or with a Pho1 gene lacking the L80 region (Pho1ΔL80). While expression of Pho1 in BMF136 restored normal wildtype phenotype, the introduction of Pho1ΔL80 enhanced growth rate and plant productivity above wildtype levels. Mass spectrometry analysis of proteins captured by anti-Pho1 showed the surprising presence of PsaC, the terminal electron acceptor/donor subunit of photosystem I (PSI). This unexpected interaction was substantiated by reciprocal immobilized protein pulldown assays of seedling extracts and supported by the presence of Pho1 on isolated PSI complexes resolved by blue native gels. Spectrophotometric studies showed that Pho1ΔL80 plants exhibited modified PSI and enhanced CO_2_ assimilation properties. Collectively, these findings indicate that the higher plant Pho1 has dual roles as a potential modulator of source and sink processes.

## Introduction

Phosphorylase is a ubiquitous enzyme (EC 2.4.1.1.) present in animals, fungi, bacteria, and plants [1]. In animals, yeast and other organisms that use glycogen as a carbohydrate reserve, this enzyme catalyzes the rate-limiting step in degradation of this α-1,4-glucan reserve by utilizing inorganic phosphate (Pi) as an acceptor and releasing Glc 1-P for its subsequent metabolism via glycolysis. The study of glycogen phosphorylase over the years has been an exquisite model for hormonal regulation of metabolism and, catalytic regulation by allosterism and reversible phosphorylation [2]. These exceptional advancements were recognized by The Nobel Foundation on two separate occasions; the Physiology or Medicine prize was awarded to Carl and Gerti Cori in 1947 [3], and later to Edmond Fischer and Edwin Krebs in 1992 [4, 5].

Despite their presence in plants, our understanding of phosphorylase’s role in plant metabolism is limited especially when compared to the wealth of knowledge on glycogen phosphorylases. In higher plants, two phosphorylase isoforms exist: the plastid-localized Pho1 (PhoL) and the cytosolic Pho2 (PhoH) [6, 7]. Both plant phosphorylases share considerable structural homology to the glycogen phosphorylases, although no evidence to date has shown that the plant enzymes are subjected to regulation by allosterism or reversible phosphorylation. The two enzyme activities have distinct roles in carbohydrate metabolism. The cytosolic Pho2 is essential for the metabolism of maltose, the amylolytic degradation product of starch that is exported from the plastid to the cytosol [8]. Based on the estimated high amounts of intracellular inorganic Pi relative to Glc 1-P levels, the plastidial Pho1 was assumed to be essential for starch turnover similar to the degradative role of glycogen phosphorylase. Such an assumption, however, was not supported by molecular studies where down-regulation of Pho1 activity (PHS1) in the leaves of *Arabidopsis thaliana* [9] and potato [7] had little influence on transitory starch metabolism, suggesting that this enzyme is not an essential component of leaf starch metabolism. Some involvement in transitory leaf starch degradation was inferred by the study of *Arabidopsis thaliana* plant lines containing mutations in PHS1 and a second gene required for metabolism of maltose, a major product of starch turnover during the night [10].

While Pho1 may be dispensable in starch degradation, substantial evidence is available that it has a major role in starch biosynthesis. This is readily evident in developing cereal grains where Pho1 expression correlates with the synthesis and accumulation of starch [11–16]. Genetic studies in the green algal *Chlamydomonas reinhardtii* [17] and the higher plant rice [11] demonstrated its substantial contribution in starch biosynthesis. Mutations in the rice Pho1 [11] or algal PhoB [17] gene resulted in significantly lower starch accumulation in both organisms, and the little amount of starch that formed had an abnormal amylopectin structure [11, 17]. In rice, the influence of Pho1 on the grain starch levels was temperature-dependent [11]. At normal growth temperature of 28°C, loss of Pho1 resulted in the formation of grains ranging in size from shrunken to pseudo-normal/white core, the latter having reduced grain weights of 90% of wildtype. In plants grown at 20°C, however, the majority of the grains were shrunken or severely shrunken. These grain phenotypes are consistent with the premise that Pho1 participates in both starch initiation and maturation of the starch grain. A failure during starch initiation causes the formation of shrunken grains, while its absence during starch grain maturation causes a reduction in the weight of mature grains. The temperature dependence of the grain phenotype suggests that Pho1 is essential for starch initiation at low temperature, but its activity is compensated at normal temperatures by an unidentified factor [11]. In addition to its impact on starch synthesis in developing grains, the *pho1^−^* rice mutant line, BMF136, matures slower as evident by the smaller vegetative biomass than normal and the delay in flowering and reproduction [11]. Hence, Pho1 activity also influences plant growth and development in general.

A biosynthetic role for Pho1 is supported by its kinetic properties. The enzyme exhibits higher catalytic efficiency in the synthesis direction with equilibrium constants ranging from 13 to 45 depending on the type of 1,4-linked α-glucan substrate tested. The synthetic capability is readily apparent when the enzyme is incubated with maltohexaose (α1,4-linked Glc_6_) in the presence of Pi, conditions favoring degradation [18]. Although the recombinant enzyme degraded Glc_6_ to smaller Glc_4_ and Glc_5_ products, it readily utilized the released Glc 1-P product and extended maltohexaose to produce larger malto-oligosaccharides at nearly the same proportion as the degradation products.

While a biosynthetic role for Pho1 in starch initiation is well established, its exact role remains unclear. The recombinant barley Pho1 was shown to *de novo* synthesize short-chain α-glucan chains in the presence of only Glc1P, indicating that the enzyme is capable of self-priming, *i.e*. does not require an existing α-1,4-linked glucan as a primer [19]. This view was not supported by a more stringent study by Nakamura *et al*. [20] where the substrate Glc 1-P was pre-treated with glucoamylase to remove any potential glucan primers [20]. Under these conditions, the rice Pho1 was incapable of self-priming starch synthesis with maltose (Glc_2_) being the smallest primer that Pho1 could extend [20].

A distinctive structural feature of the higher plant Pho1 is the presence of a 50 to 82 amino acid sequence long region (termed L80 in this study) with unknown function located about in the middle of the primary sequences of various plant species [1]. The L80 region is unique to the higher plant Pho1 and is absent in animal and fungi glycogen phosphorylases as well as in the higher plant Pho2 or in the *Chlamydomonas* phosphorylase enzymes [17].

Molecular modeling of the rice Pho1 indicates that the L80 is an intrinsically disordered peptide, structurally independent from the rest of the protein. Consistent with the structural model prediction, removal of L80 peptide (Pho1ΔL80) has no significant effect on the kinetic properties of recombinant rice Pho1 and, therefore, is not required for the catalysis [18, 21].

To elucidate the role of the Pho1 L80 region, we complemented the *pho1^−^* BMF136 with wildtype Pho1 or Pho1ΔL80 gene sequences. Complementation of BMF136 with wildtype Pho1 restored grain weight and growth properties to their normal levels. Surprisingly, expression of Pho1ΔL80 enhanced grain weight and growth properties above wildtype levels. Protein-protein interaction studies showed that Pho1 or Pho1ΔL80 interacts with the PsaC subunit of photosystem I (PSI). While Pho1 plants exhibited PSI properties similar to wildtype, Pho1ΔL80 (Pho1ΔL80) showed significantly altered PSI properties and enhanced CO_2_ assimilation rates. Overall, these results indicate that the higher plant Pho1 has two distinct roles in starch biosynthesis and, unexpectedly, in modulating PSI activity.

## Results

### Introduction of Pho1 to BMF136 restores wildtype grain weight while the introduction of Pho1ΔL80 elevates it above wildtype levels

To gain insight on the function of the non-catalytic L80-peptide domain, the *pho1^−^* deficient BMF136 rice line was transformed with wildtype Pho1 and Pho1ΔL80 gene sequences. Expression of these two Pho1 cDNA sequences was directed by a near intact, native Pho1 promoter, which was missing a portion of the TC-repeat region located at the 3’promoter end. These constructs also contained the Rubisco small subunit transit peptide to direct the newly synthesized Pho1 to the plastid **(Figure 1A)**. A similar pair of Pho1 and Pho1ΔL80 gene sequences were also constructed with the stronger AGPS2 promoter for overexpression **(Supplementary Figure 1A)**, the latter exhibiting a temporal mRNA expression similar to the Pho1 gene [12]. The amino-terminal part of the Pho1 cDNA was replaced by Pho1 genomic sequence containing plastid targeting sequence and first intron **(Supplementary Figure 1A)**.

**Figure 1:**
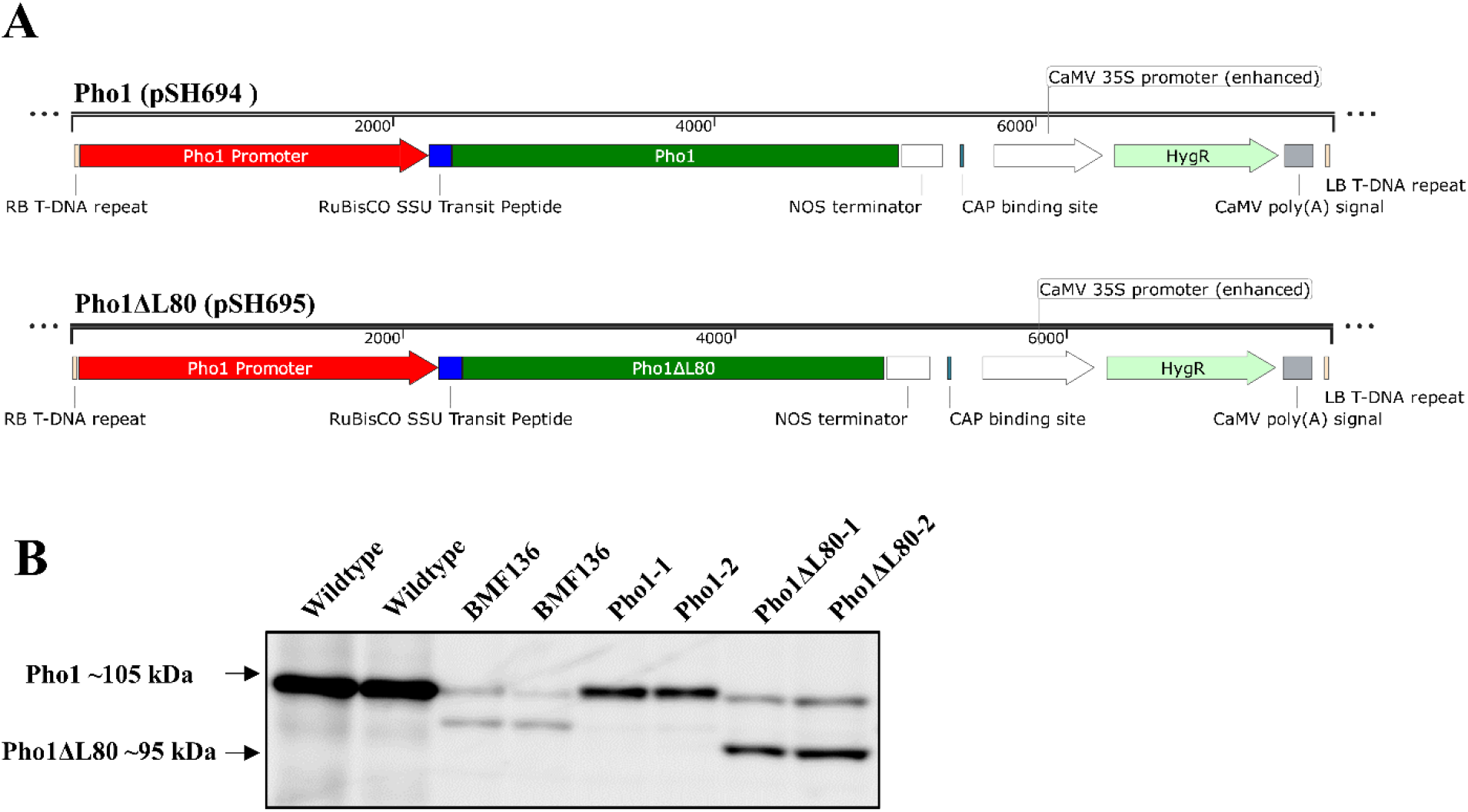
Expression of Pho1 or Pho1ΔL80 in Pho1-deficient BMF136. (A) Maps (created by SnapGene) of DNA constructs transformed into BMF136. (B) Representative expression levels of Pho1 and Pho1ΔL80 in mature seeds of transgenic rice lines. One seed from each plant was crushed and proteins extracted using 200 μL of Urea/SDS extraction buffer. 7.5 μL of the sample was resolved on a 7.5% SDS polyacrylamide gel and then transferred onto a nitrocellulose membrane. Anti-Pho1 antibody was used to examine the relative expression levels of Pho1 (105 kDa) and Pho1ΔL80 (95 kDa) for each plant. Note that BMF 136 appears to express low levels of wildtype Pho1 as well as variant form lacking 56 residues in exon 5.

Independent transformation events were selected using hygromycin resistance, propagated, and screened. The first two homozygous lines identified for each construct were used in this study. Expression of Pho1 or Pho1ΔL80 in *pho1^−^* deficient BMF136 and their relative levels in the parental line BMF136 and wildtype (TC65) were evaluated by immunoblot analysis of mature grain extracts. Production of Pho1 or Pho1ΔL80 protein was restored in the BMF136 background, but to different extents (**Figure 1B**). Pho1 or Pho1ΔL80 protein levels were lower in Pho1 or Pho1ΔL80 rice lines employing the native Pho1 promoter sequences than wildtype, while expression of Pho1 and Pho1ΔL80 in overexpression lines employing the AGPS2 promoter were higher than wildtype (**Supplementary Figure 1B**).

BMF 136 exhibited two faint polypeptide bands, one at the same size as Pho1 and a second smaller one. BMF136 was previously shown to have a GT to AT mutation at the 5’ splice site of the 5^th^ intron in the Pho1 gene, which prevented removal of the 5^th^ intron and nonsense-mediated mRNA decay [11]. Additional cDNA sequence analysis showed, however, that an AG-GT 5’splice site within the 5^th^ exon was used instead to produce a smaller variant protein lacking 53 amino acids of exon 5 **(Figure S2)**. Thus, it appears that the lower polypeptide band corresponds to this smaller variant Pho1 subunit. The upper band corresponds to the size of the wildtype Pho1 and was likely generated by weak splicing activity of the 5^th^ intron. These results indicate that BMF136 is not a null *pho1^−^* mutant but may possess very low level of intact Pho1 protein and, in turn, catalytic activity.

Interestingly, the signals from the weakly spliced Pho1 and smaller Pho1 variant differed in the transgenic lines compared to BMF136. The weakly spliced Pho1 wildtype band was slightly enhanced in the Pho1ΔL80 transgenic lines, while the smaller Pho1 variant protein band detected in BMF136 was significantly reduced or absent in the Pho1 and Pho1ΔL80 transgenic lines. The basis for the enhancement of the weakly splice variant polypeptide band in Pho1ΔL80 plants is likely due to its increased stability by its forming a dimer with Pho1ΔL80 protein. It is not clear why the amounts of the small Pho1 variant are reduced or absent in the Pho1ΔL80 transgenic lines.

### Pho1ΔL80 plants produce larger grains and exhibit faster growth rates than the wildtype

As noted earlier [11], the BMF136 mutant line exhibits a grain weight distribution ranging from severely shrunken to normal-appearing grains (pseudo-normal) **(Figure 2B)**. To determine if the grain weight distribution pattern of BMF136 was restored to a wildtype pattern in transgenic lines, weights of individual grains from wildtype, BMF136 and transgenic lines were measured. Complementation of BMF136 plants with the Pho1gene restored both grain weight and grain weight distribution at levels comparable to wildtype (**Table 1 and Figure 2A, B, C and D**). Unexpectedly, BMF136 plants expressing the Pho1ΔL80 gene produced grains with increased weight (~27% above wildtype levels) and increased grain yields (~33% above wildtype levels) (**Table 1**). The grain weight distribution of Pho1ΔL80 lines was also displaced to a higher range than that observed for wildtype (**Figure 2E and F).** These results indicate that even though protein expression levels of Pho1 or Pho1ΔL80 in Pho1 rice lines were lower than in wildtype, they were sufficient to restore normal grain weight and grain weight distribution. Therefore, only a portion of the amount of Pho1 activity in wildtype plant is sufficient for normal starch synthesis during grain development.

**Figure 2:**
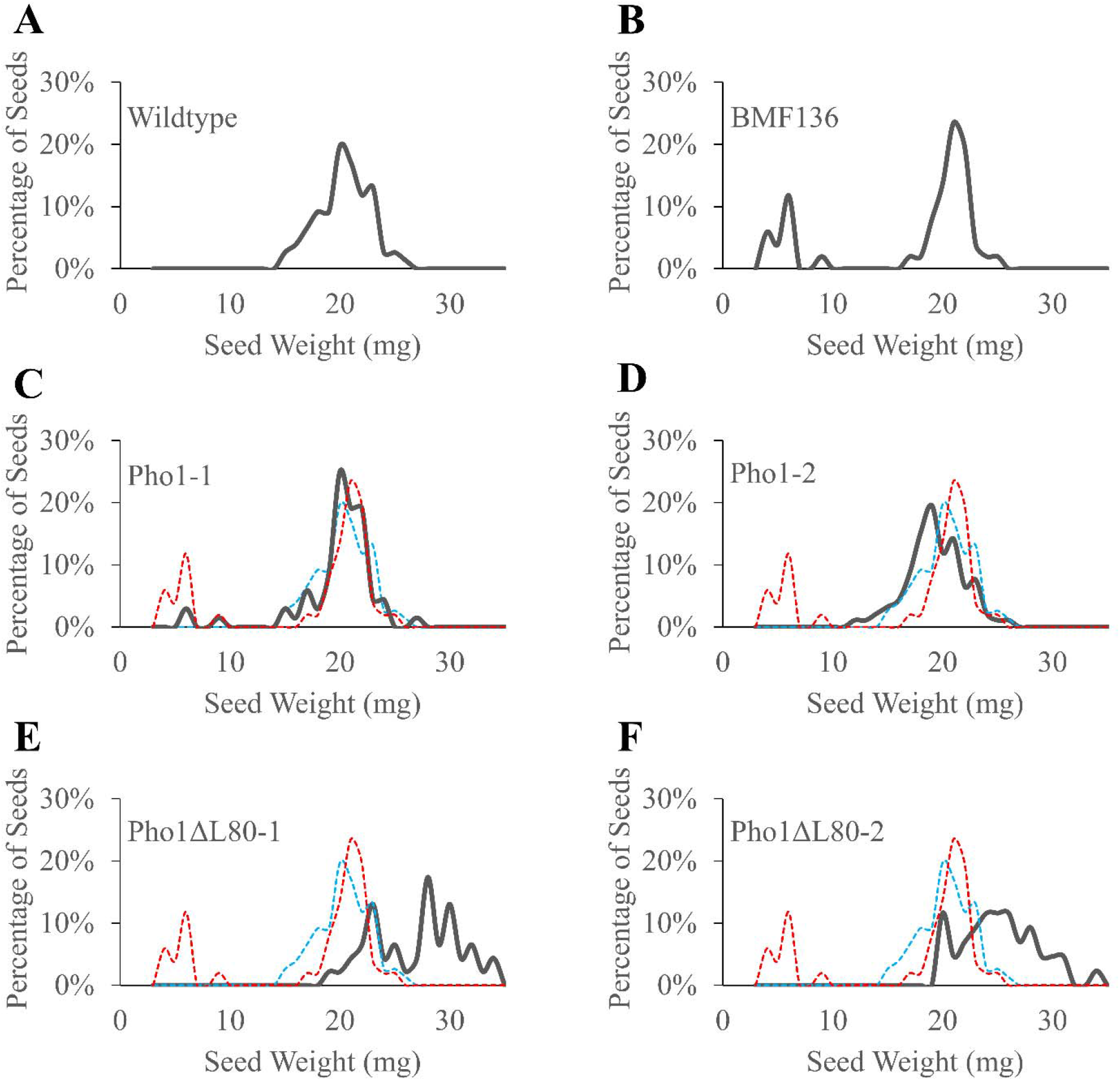
Seed weight distribution of (A) wildtype, (B) BMF136, (C-D) Pho1 and (E-F) Pho1ΔL80 plants. Bold black lines show the percentage of seeds with a given weight harvested from the indicated genotype. Faint blue and red dashed lines show the seed weight distribution of wildtype and BMF136 plants, respectively, for comparison with the transgenic mutants. Seed weight distribution was determined by weighing individual seeds pooled from four sibling plants from the given line grown in parallel.

**Table 1:**
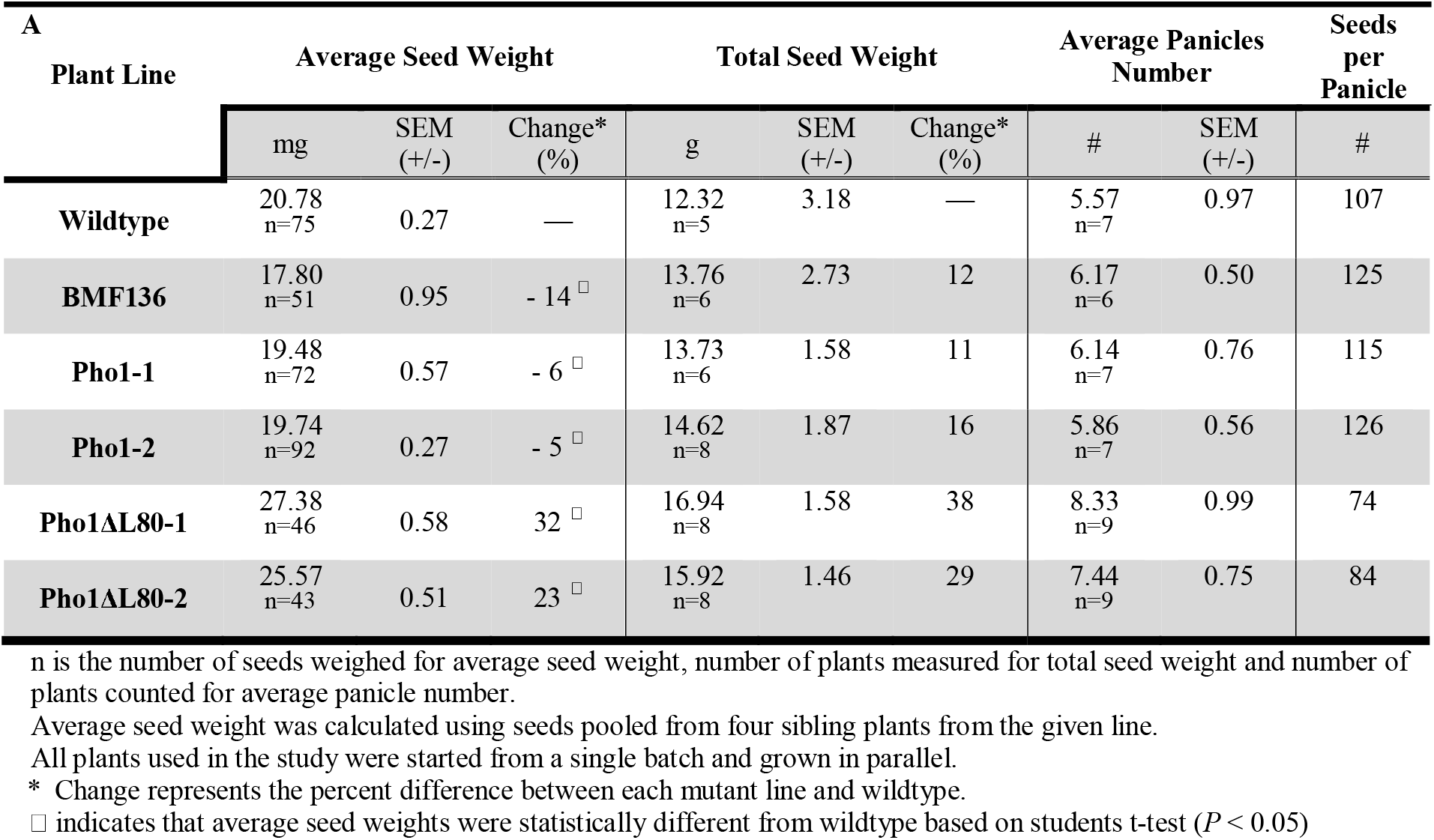
Seed weight characteristics of wildtype, BMF mutant, and transgenic rice plants. Note that BMF plants were harvested later than the other lines, due to delayed maturation, which accounts for the higher total seed weight and average panicle number.

| A<br>Plant Line | Average Seed Weight |  |  | Total Seed Weight |  |  | Average Panicles Number |  | Seeds per Panicle |
| --- | --- | --- | --- | --- | --- | --- | --- | --- | --- |
|  | mg | SEM (+/-) | Change* (%) | g | SEM (+/-) | Change* (%) | # | SEM (+/-) | # |
| **Wildtype** | 20.78<br>n=75 | 0.27 | — | 12.32<br>n=5 | 3.18 | — | 5.57<br>n=7 | 0.97 | 107 |
| **BMF136** | 17.80<br>n=51 | 0.95 | - 14 □ | 13.76<br>n=6 | 2.73 | 12 | 6.17<br>n=6 | 0.50 | 125 |
| **Pho1-1** | 19.48<br>n=72 | 0.57 | - 6 □ | 13.73<br>n=6 | 1.58 | 11 | 6.14<br>n=7 | 0.76 | 115 |
| **Pho1-2** | 19.74<br>n=92 | 0.27 | - 5 □ | 14.62<br>n=8 | 1.87 | 16 | 5.86<br>n=7 | 0.56 | 126 |
| **Pho1ΔL80-1** | 27.38<br>n=46 | 0.58 | 32 □ | 16.94<br>n=8 | 1.58 | 38 | 8.33<br>n=9 | 0.99 | 74 |
| **Pho1ΔL80-2** | 25.57<br>n=43 | 0.51 | 23 □ | 15.92<br>n=8 | 1.46 | 29 | 7.44<br>n=9 | 0.75 | 84 |

n is the number of seeds weighed for average seed weight, number of plants measured for total seed weight and number of plants counted for average panicle number.
Average seed weight was calculated using seeds pooled from four sibling plants from the given line.
All plants used in the study were started from a single batch and grown in parallel.
\* Change represents the percent difference between each mutant line and wildtype.
□ indicates that average seed weights were statistically different from wildtype based on students t-test ( $P < 0.05$ )

These various rice lines exhibited significantly different reproductive properties. While wildtype plants harbored more grains per panicle than Pho1ΔL80, the Pho1ΔL80 plants contained more panicles per plant than wildtype **(Table 1)**. Hence, the total number of grains produced by these two plant lines were similar. The higher grain biomass produced by the Pho1ΔL80 transgenic lines is due to the larger grain size than those from wildtype plants.

The BMF136 mutant line produced smaller grains on average. Unlike wildtype and the Pho1/Pho1ΔL80 rice lines, whose vegetative growth essentially stopped at the flowering stage, BMF136 continued to grow and produced additional tillers due to delayed maturation. Hence, total grain weights and grains per panicle values were higher for BMF136 **(Table 1)** than the other lines due to the much longer growth period.

Pho1ΔL80 plants grew faster than wildtype. This is readily seen in the increase panicle number produced by Pho1ΔL80 transgenic lines compared to wildtype when these plants were harvested at the same time. To study the growth rates in detail, seedlings of the various rice lines were grown in parallel and plant heights compared at 7, 14 and 21 days after germination (DAG) (**Figure 3**). While wildtype, BMF136 and Pho1 lines showed similar growth rates (Figure 3, **Table 2, control column),** those for **the** Pho1ΔL80 transgenic lines were significantly higher even at 7 DAG.

**Figure 3:**
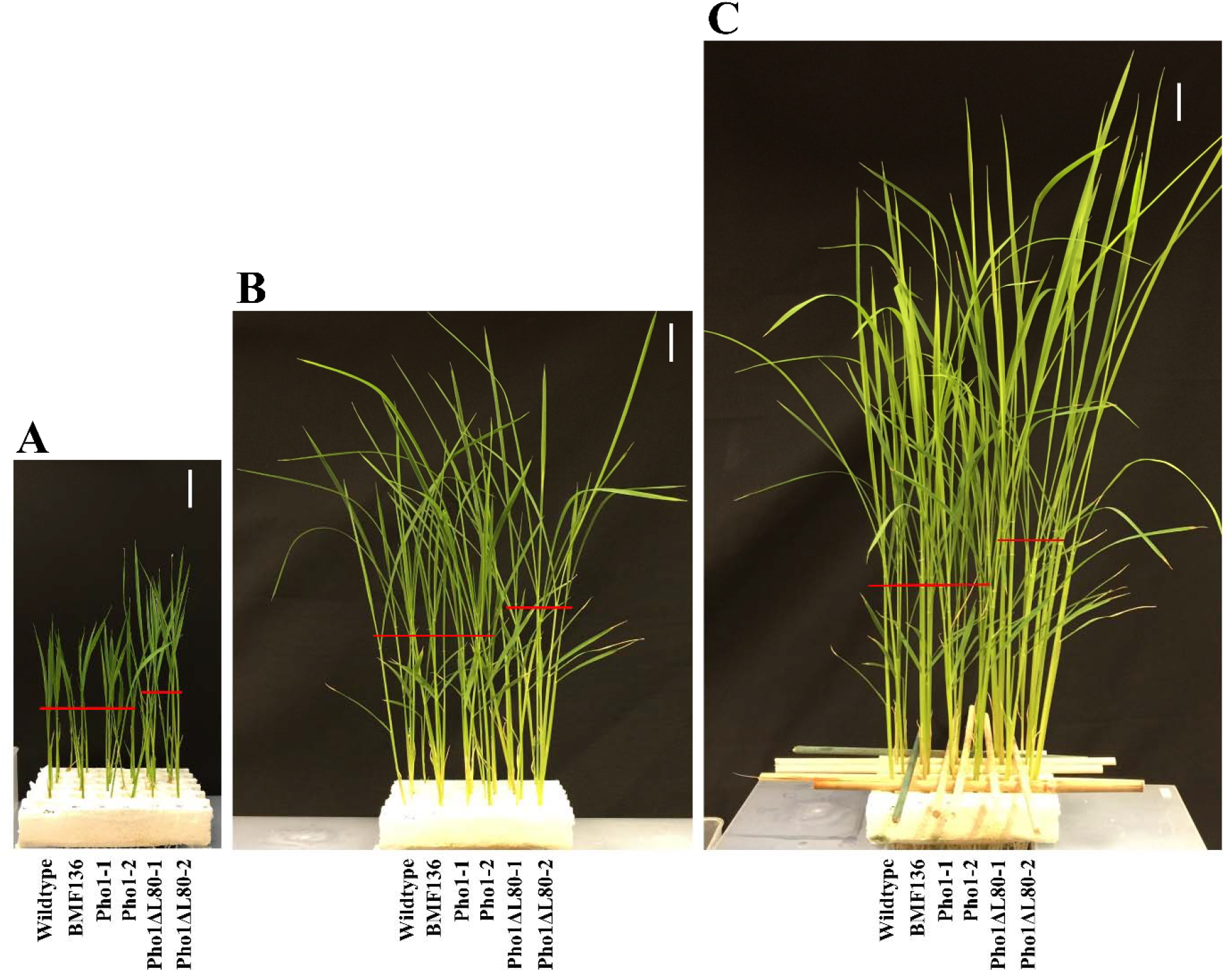
Growth rates of wildtype, BMF136, Pho1 and Pho1ΔL80 plants grown in parallel. (A) 7 DAG, (B) 14 DAG and (C) 21 DAG. Red lines represent the proximate height of the top node. White scale = 2.5 cm. DAG = days after germination.

**Table 2:** Heights of wildtype, BMF mutant, and transgenic rice plants under control, phosphorus deficient or nitrogen deficient conditions at 21 DAG.

| Plant Line | Control |  |  | Phosphorus Deficient |  |  | Nitrogen Deficient |  |  |
| --- | --- | --- | --- | --- | --- | --- | --- | --- | --- |
|  | Height (cm) | SEM (+/-) | Change* (%) | Height (cm) | SEM (+/-) | Change* (%) | Height (cm) | SEM (+/-) | Change* (%) |
| **Wildtype** | 12.98<br>n=5 | 0.47 | — | 12.22<br>n=5 | 0.34 | — | 7.88<br>n=5 | 0.51 | — |
| **BMF136** | 12.42<br>n=5 | 0.69 | - 4.3 | 12.16<br>n=5 | 0.36 | - 0.5 | 7.68<br>n=5 | 0.25 | - 2.5 |
| **Pho1-1** | 12.76<br>n=5 | 0.55 | - 1.7 | 11.72<br>n=5 | 0.31 | - 4.1 | 7.56<br>n=5 | 0.62 | - 4.1 |
| **Pho1-2** | 13.50<br>n=4 | 0.31 | 4.0 | 11.64<br>n=5 | 0.47 | - 4.7 | 7.90<br>n=5 | 0.26 | 0.3 |
| **Pho1Δ80-1** | 15.38<br>n=5 | 0.94 | 18.5 | 14.50<br>n=5 | 0.28 | 18.7 □ | 10.54<br>n=5 | 0.62 | 33.8 □ |
| **Pho1Δ80-2** | 15.62<br>n=5 | 0.43 | 20.3 □ | 14.36<br>n=5 | 0.78 | 17.5 | 10.90<br>n=5 | 0.78 | 38.3 □ |

\* Change represents the percent difference between each mutant line and wildtype under tested condition.
□ indicates that average seed weights were statistically different from wildtype based on students t-test ( $P < 0.05$ )
All plants used in the study were started from a single batch and grown in parallel.

### Pho1 interacts with the PsaC subunit of PSI

The restoration of average grain weight and grain weight distribution by expression of wildtype Pho1 in BMF136 supports Pho1’s dual role in starch initiation and starch grain maturation. The larger grains produced by Pho1ΔL80 lines indicate that starch production is elevated during grain development. Pho1ΔL80 rice lines also exhibited higher growth rates than wildtype plants. These results indicate that the L80 peptide domain acts as a negative modulator of starch synthesis and growth. The effect on growth was thought to be an indirect result of starch metabolism stimulating photosynthesis [22, 23]. The following results, however, support a direct effect of Pho1 on photosynthesis and, specifically, PSI.

To gain insight on how Pho1 may influence the growth rates and seed biomass production in rice plants, proteomic studies were initiated. In two independent experiments, developing seed extracts were incubated with immobilized affinity-purified anti-Pho1 and the captured proteins analyzed by SDS polyacrylamide gel electrophoresis and LC-MS/MS (**Figure 4**). Compared to the IgG control, the Pho1 antibody captured Pho1 and a diverse array of polypeptides as resolved by silver-stained polyacrylamide gel electrophoresis (**Figure 4A**). Subsequent proteomic analysis of the immunoprecipitated proteins from each experiments revealed the presence of two high confidence unique peptides belonging to photosystem-I subunit C (PsaC). These two peptides accounted for 32% of the full-length PsaC protein. In addition to PsaC, analysis of the two trials identified twenty other proteins captured by anti-PhoI and not by IgG control. In addition to Pho1, 4-α-glucantransferase I (Dpe1) was also found in anti-Pho1 immunoprecipitates (**Figure 4B, Supplementary Data**). The presence of Dpe1 in the anti-Pho1 IP was expected as this protein is associated with Pho1 as a protein complex in rice seed extracts [21]. Interestingly, starch branching enzymes, which have been established to interact with Pho1 in rice endosperm [20], were not detected.

**Figure 4:**
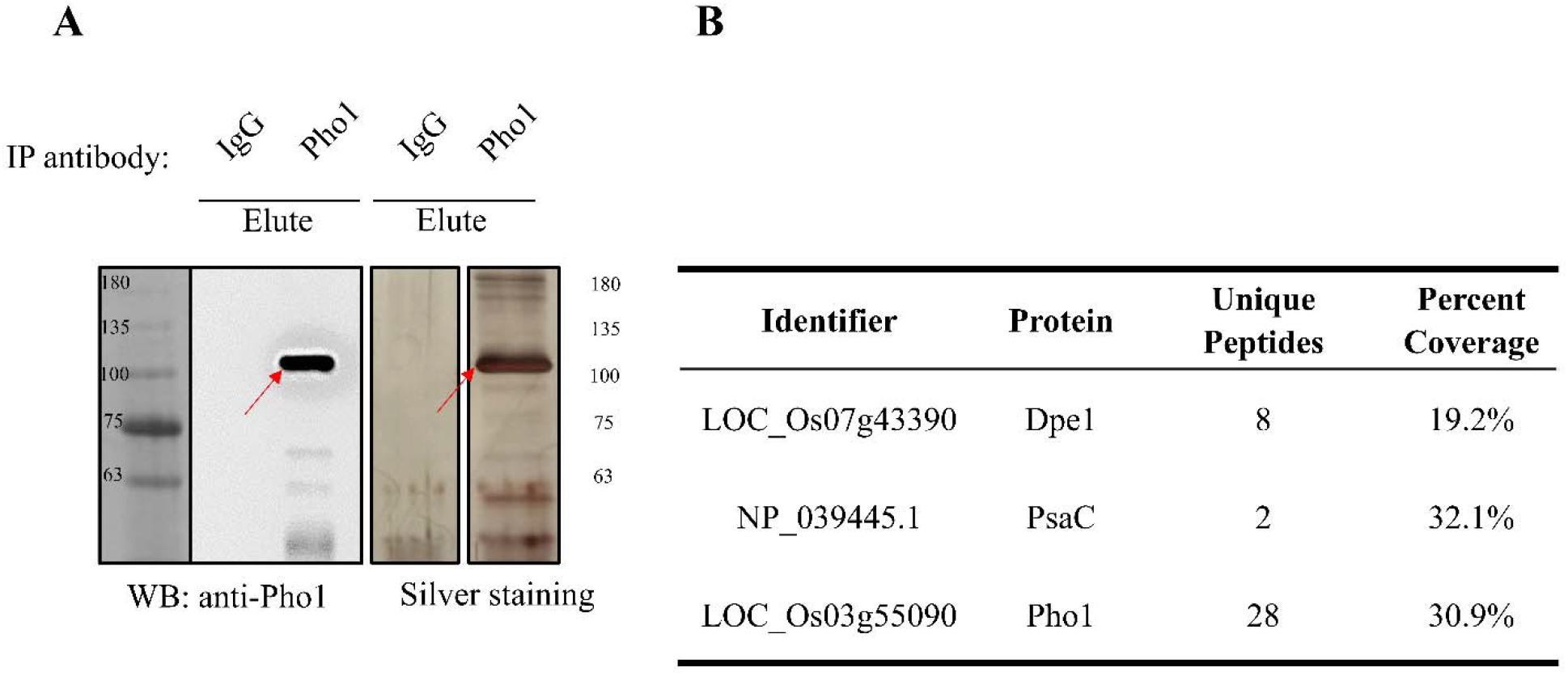
Identification of proteins co-immunoprecipitated with Pho1. (A) Immunoblot analysis and silver staining of eluted proteins after immunoprecipitation of Pho1 using anti-Pho1 antibody. The red arrows indicate immunoprecipitated 105 kDa Pho1. Rabbit Immunoglobulin G (IgG) is used as the antibody control. 10 μL of 100 μL total IP eluate was loaded to each well. (B) Prominent co-immunoprecipitated proteins identified by LC/MS/MS.

The interaction of Pho1 with the photosynthetic-related PsaC was unexpected, but could account for the differences in growth between BMF136, Pho1 and Pho1ΔL80 lines. To obtain further evidence for a relationship between these two proteins, we first studied the expression patterns of Pho1 and PsaC in leaves, mid-developing grains, sheaths, and seedlings (7-10 DAG) by immunoblot analysis (**Figure 5A**). Both Pho1 and PsaC were detected in all four tissues although the highest co-expression was evident in seedlings (minus root tissue).

**Figure 5:**
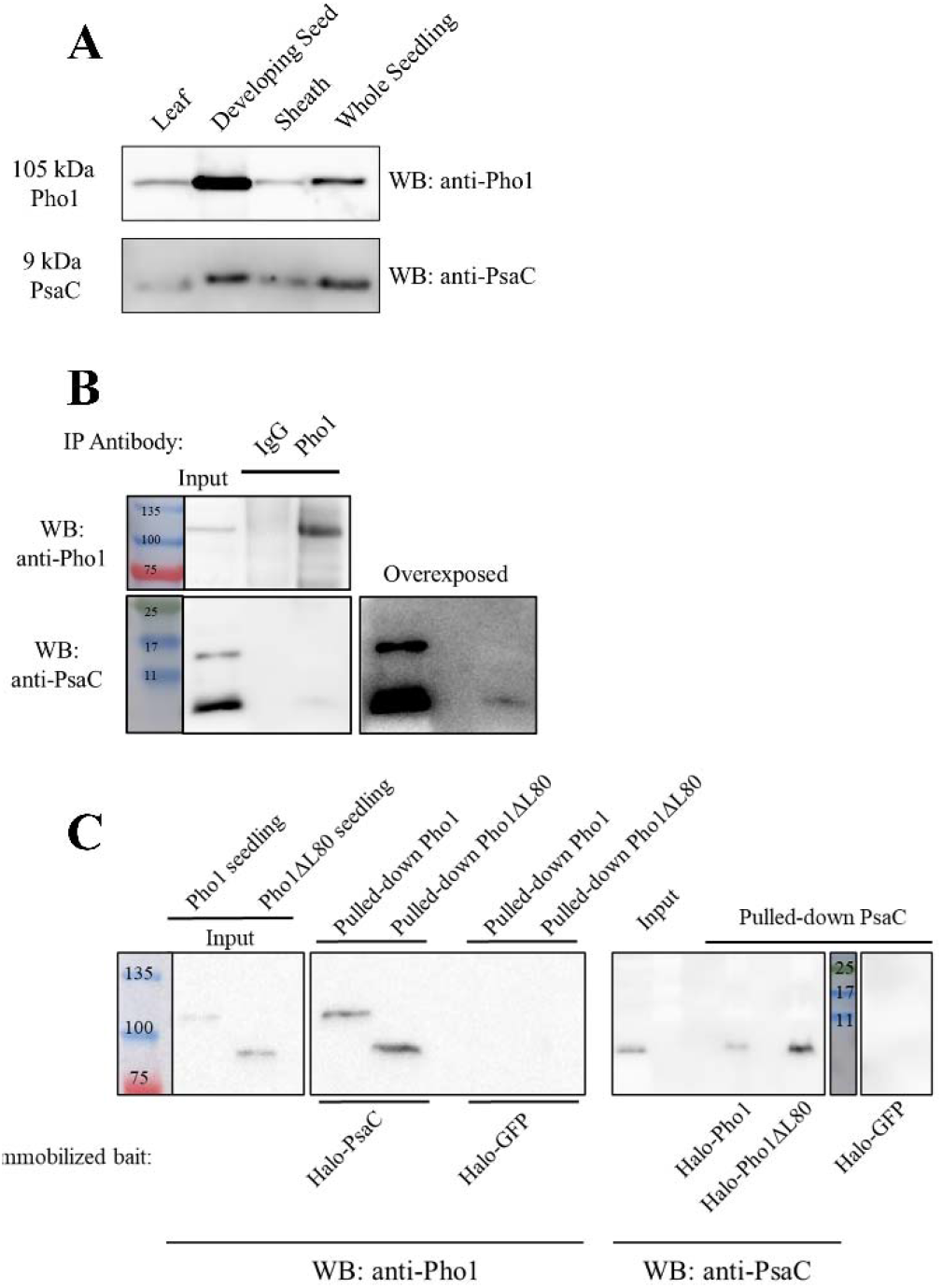
Verification of the interaction between Pho1 and PsaC. (A) Expression of 105 kDa Pho1 and 9 kDa PsaC in leaves, developing seeds, sheaths and seedlings. Each lane contained 250 μg of total tissue for detecting Pho1 and 750 μg of total tissue for detecting PsaC, except the PsaC leaf sample containing 75 μg of total tissue (B) Coimmunoprecipitation of PsaC when seedling green tissue extracts were incubated with anti-Pho1. Upper panel depicts immunoblot probing with anti-Pho1 while the lower panel was probed with anti-PsaC. The very right panel depicts an over-exposed image to show the low but detectable amounts of PsaC present in the Co-IP. (C) The capture of Pho1 or Pho1ΔL80 from 60 μg of AGPS2-Pho1 and AGPS2-Pho1ΔL80 green seedling extracts by immobilized Halo-PsaC (left panel) and the capture of PsaC from 60 μg of wildtype green seedlings by using immobilized Halo-Pho1 or Halo-Pho1ΔL80 (right panel).

Using seedling extracts as a source, anti-Pho1 readily captured PsaC (**Figure 5B**). Due to the relatively low titer of the commercially available anti-PsaC antibody, a reciprocal PsaC Co-IP could not be conducted. Thus, to validate the interaction between Pho1 and PsaC, reciprocal pulldown assays employing immobilized Halo-Pho1, Halo-Pho1ΔL80, and Halo-PsaC were used to further substantiate the interaction between Pho1 and PsaC. For Halo-PsaC pulldowns, seedling extracts were prepared from A_p_Pho1 and ApPho1ΔL80 plants due to the higher expression of transgenic Pho1 or Pho1ΔL80 proteins in these lines. Figure 5C readily shows that immobilized PsaC captured native Pho1 and Pho1ΔL80 while both immobilized Pho1 and Pho1ΔL80 captured native PsaC. Although reciprocal interactions were readily evident between PsaC and Pho1 or between PsaC and Pho1ΔL80, PsaC did not interact equally with Pho1 and Pho1ΔL80 as the PSI subunit preferred the latter (see right panel, **Figure 5C**).

Pho1 [11] and PsaC [24] are plastid-localized proteins. While Pho1 is encoded by the nuclear genome and transported to the plastid stroma, PsaC, on the other hand, is encoded by the chloroplast genome [25]. PsaC is a soluble protein, and it is bound to PSI facing the stroma side of the thylakoid membrane [24]. To test whether Pho1 was associated with PSI, intact thylakoid membranes were isolated from rice seedling chloroplasts and solubilized PSI and other protein complexes resolved by blue-native PAGE (**Figure 6A**). PSI containing complexes were identified based on previous reports [26–29] and determined to correspond to bands I (PSI-NDH super-complex, >1000 kDa), II (PSI + LHCI, 568 kDa) and III (PSI, 379 kDa) (**Figure 6A**). Proteins from these bands were gel extracted, denatured, dot-blotted and probed with anti-Pho1. Bands IV and V, which were devoid of PSI, were used as internal controls. As seen in **Figure 6B**, Pho1 signal was detected in PSI containing bands I, II and III but not in the non-PSI control bands, IV and V.

**Figure 6:**
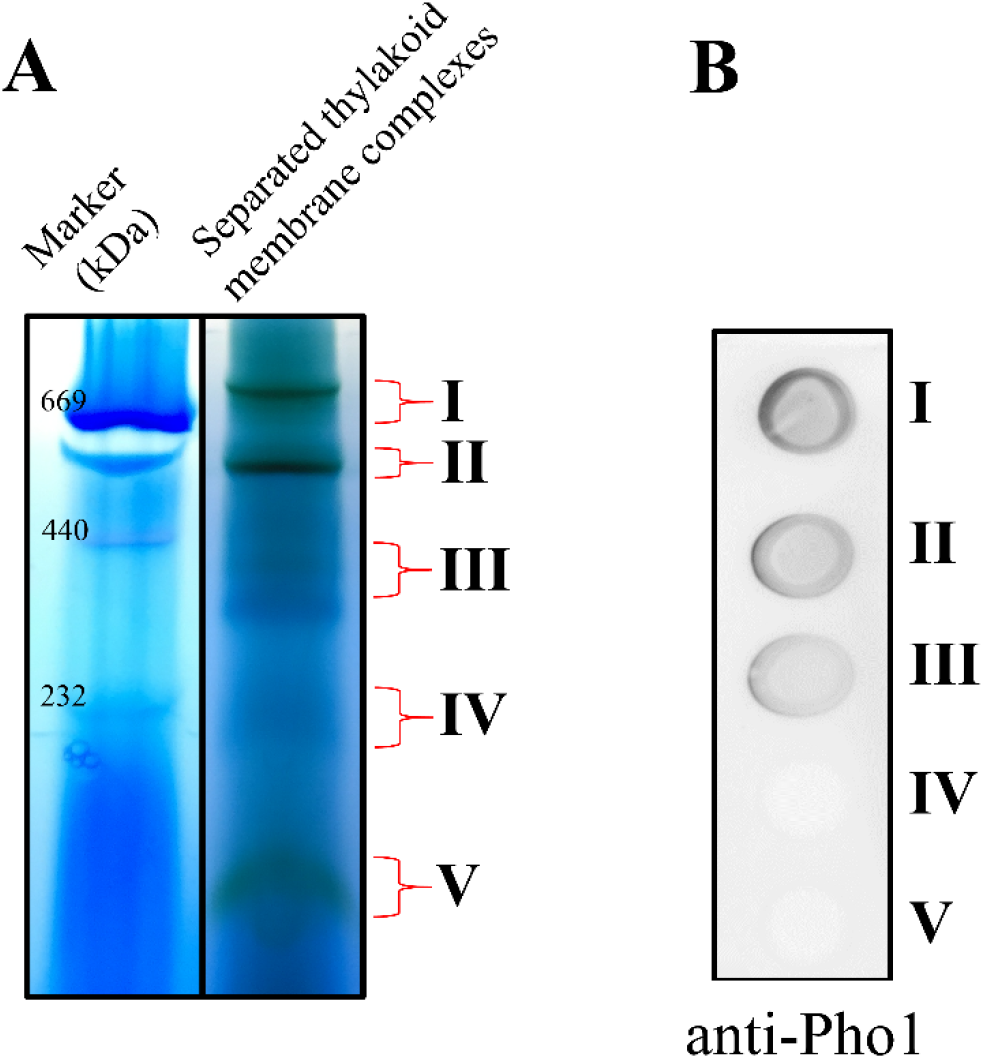
Identification of the sub-chloroplastic location of Pho1-PsaC interaction. (A) Thylakoid membranes were isolated from purified chloroplasts and protein complexes separated by blue-native PAGE. PSI is detected in bands I (PSI-NDH super-complex, >1000 kDa), II (PSI + LHCI, ~ 550 kDa) and III (PSI, ~ 400 kDa). Bands IV and V lack PSI. (B) Dot-blot analysis for the presence of Pho1 in the bands I to V from blue-native gel.

### Pho1ΔL80 mutants exhibit altered PSI parameters

To assess whether the interaction between Pho1 and PsaC has physiological significance, electron transport properties of PSI of the various rice lines grown in parallel were evaluated using a flash-type spectrophotometer that probes the light-induced redox changes of the reaction center chlorophyll P700 in intact plants based on difference absorption measurements at 820 nm and 900 nm [30]. From these measurements three PSI parameters were determined as function of light intensities [30, 31]. The first parameter (YI) is the quantum yield of photochemical energy conversion of PSI, the second (YN_D_), is the non-photochemical quantum yield due to PSI donor side limitations, *i.e*. electron transport from cyt b_6_f via plastocyanin, while the third parameter (YN_A_) is the non-photochemical quantum yield due to PSI acceptor side limitations, *i.e*. electron transport from PSI to ferredoxin and other downstream acceptors. These three quantum yield values were determined for wildtype, BMF136, Pho1 and Pho1ΔL80 plants (28-35 days after transplantation, DAT).

The overall trend of the YI, YN_D_, and YN_A_ parameters as function of light intensity (**Figure 7**) was observed as expected [30, 31]. YN_A_ declines with light intensity due to activation of electron sinks, mainly Calvin-Benson-Bassham cycle. In contrast, the donor side of PSI (YN_D_) becomes more limiting at higher light intensities due to the stronger build-up of a proton gradient across the thylakoid membrane in the light as electron flow through the cytochrome b_6_f slows down (photosynthetic control). Finally, the photochemical energy conversion efficiency of PSI (YI) declines with increasing light intensities mainly as a consequence of the increase PSI donor side limitation (YN_D_) [30].

**Figure 7:**
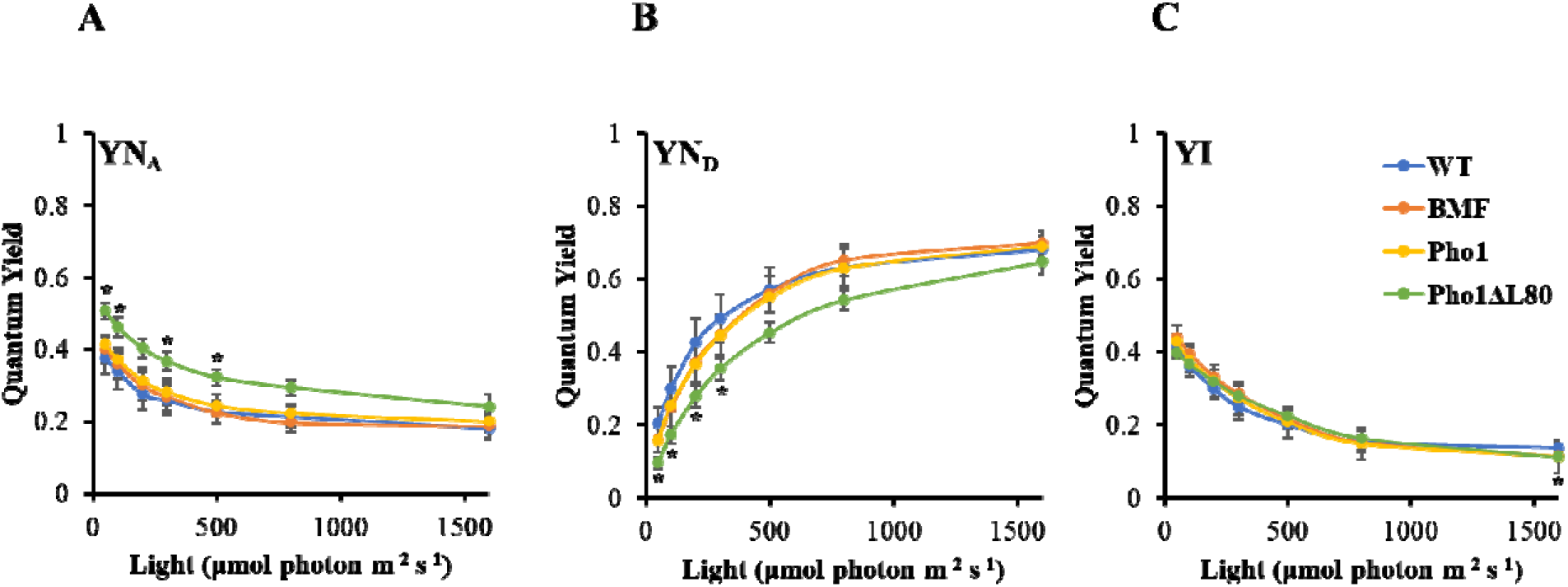
PSI properties of wildtype, BMF136, Pho1 and Pho1ΔL80 plants investigated through P700 redox state measurements. (A) Non-photochemical quantum yield of the acceptor site limitation, YN_A_ (+/- SEM) in response to light intensity. (B) Non-photochemical quantum yield of the donor site limitation, YN_D_ (+/- SEM), in response to light intensity. (C) Quantum yield of the photochemical energy conversion, YI (+/- SEM), in response to light intensity. * p-value < 0.05 with respect to wildtype. The numbers of plants analyzed wildtype (n=12), BMF136 (n=14), Pho1 (n=29), and Pho1ΔL80 (n=32). Measurements are done by two different investigators.

Among the various plant lines, Pho1ΔL80 was the only line that displayed significant differences in the various functional PSI parameters. Pho1ΔL80 showed higher PSI acceptor side limitation (higher YN_A_) (**Figure 7A**) and a lower PSI donor side limitation (lower YN_D_) compared to the other plant lines **(Figures 7B)**. Interestingly, the increase in acceptor side limitation and the decrease in donor side limitation in Pho1ΔL80 compensated each other, leading to an unchanged YI (**Figures 7B and 7C**). Changes in PSI donor and acceptor side properties of Pho1ΔL80 were also relatively light intensity independent, *i.e*. YN_A_ and YN_D_ show a constant offset over the measured intensity range (**Figure 7A and 7B**). The alteration of PSI properties in the Pho1ΔL80 lines was not related to PSII activity as photosystem-II (PSII) quantum yields showed no difference between the various rice lines **(Figure 8C)**.

**Figure 8:**
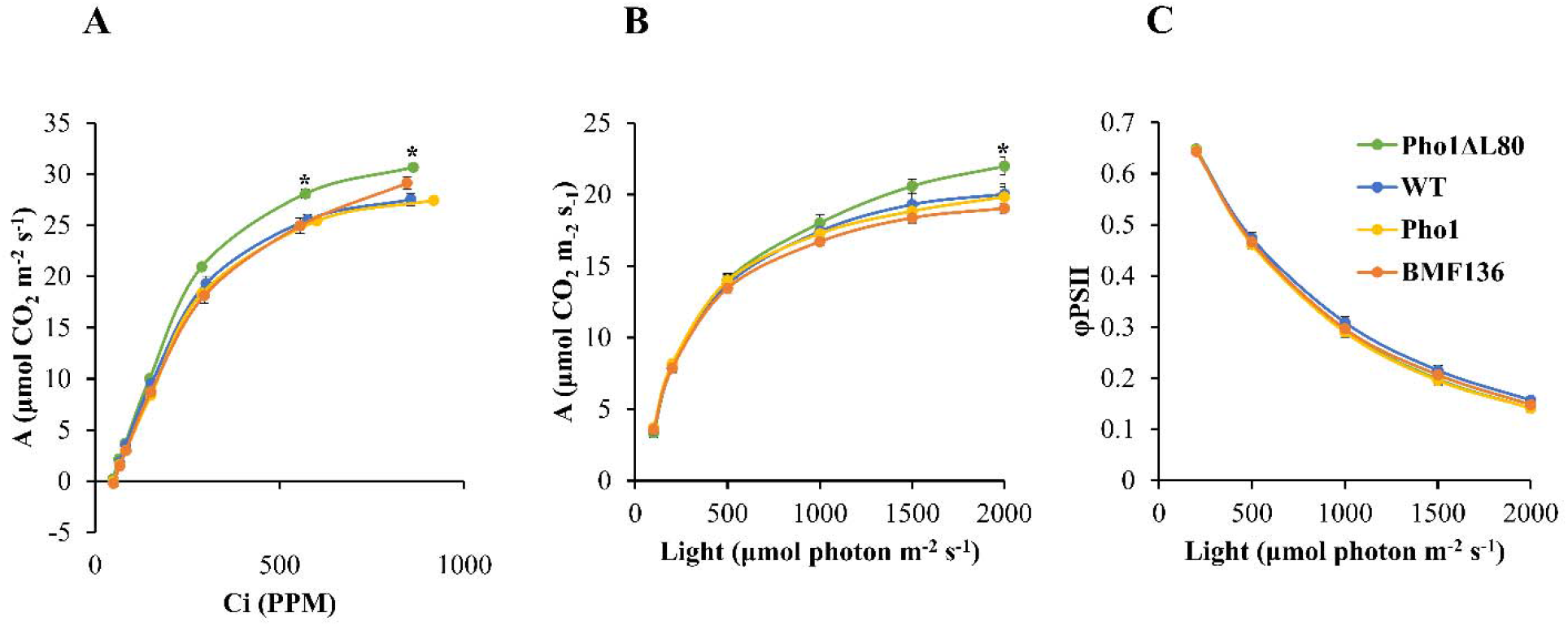
The net rates of CO_2_ assimilation and chlorophyll florescence of wildtype, BMF136, Pho1 and Pho1ΔL80 plants. (A) CO_2_ assimilation rate (+/- SEM) in response to changing intercellular CO_2_ concentrations (Ci: 50, 75,100, 200, 400, and 800 PPM) at 1500 μmol photon m^−2^ s^−1^ and 27°C. (B) CO_2_ assimilation rate (+/- SEM) in response to changing light (PAR: 0, 100, 200, 500, 1000, 1500, 2000 μmol photon m^−2^ s^−1^) at 400 PPM CO_2_ and 27°C. (C) Photosystem-II quantum yield (+/- SEM) in response to changing light (PAR: 0, 100, 200, 500, 1000, 1500, 2000 μmol photon m^−2^ s^−1^) at 400 PPM CO_2_ and 27°C. * p-value < 0.05 with respect to wildtype. Numbers of plants examined are wildtype (n=6), BMF136 (n=6), Pho1 (n=14), and Pho1ΔL80 (n=14).

### Pho1ΔL80 mutants exhibit higher CO_2_ assimilation rates

Gas exchange measurements were conducted to determine if the alterations of the PSI properties affected the capacity to fix carbon. The net rates of CO_2_ assimilation were measured for wildtype, BMF136, Pho1 and Pho1ΔL80 plants grown in parallel at 35-45 DAT under constant light intensity (1500 μmol photon m^−2^ s^−1^) with increasing CO_2_ (CO_2_ response), or under constant CO_2_ (400 ppm) with increasing light intensity (light response), (**Figure 8**).

When measured for CO_2_ response, Pho1ΔL80 lines exhibited higher net rates of CO_2_ assimilation than wildtype and the other rice lines (**Figure 8A**). In terms of light response, Pho1ΔL80 lines had higher net rates of CO_2_ assimilation that trended higher than wildtype above 1,000 μmol photons m^−2^ s^−1^ (**Figure 8B)**. However, this difference was only significant at 2,000 μmol photons m^−2^ s^−1^. While BMF136 trended lower than wildtype above 500 μmol photons m^−2^ s^−1^, the difference was not statistically significant at any point.

### Pho1ΔL80 mutants are less susceptible to nitrogen starvation

The growth patterns of transgenic plant lines were evaluated, in parallel, under control, nitrogen- or phosphorus-deficient conditions. Under control conditions, Pho1ΔL80 expressing transgenic lines were 18-20% taller than wildtype at 21 DAG. However, under nitrogen starvation, the height difference between Pho1ΔL80 expressing transgenic lines and the wildtype increased to 34-38% (Table 2). Hence, Pho1ΔL80 lines are more efficient at metabolizing nitrogen than wildtype. By contrast, there was no significant percent differences in plant heights between wildtype and Pho1 lines under normal and phosphorous-deficient conditions (Table 2).

### Pho1ΔL80 mutants are susceptible to cold stress

PSI is known to be particularly sensitive to cold stress [32]. As Pho1 may modulate PSI activity via its interaction with PsaC *in vivo*, the cold tolerance of wildtype, BMF136, Pho1 and Pho1ΔL80 plants was investigated. Plants were grown at a constant temperature of 20°C, the lowest temperature that enables pollen production and successful fertilization to occur in rice [33]. The contents of PsaB (PSI core protein) of the cold grown wildtype, BMF136, Pho1 and Pho1ΔL80 plants were estimated at 120 DAT by immunoblot analysis of leaf extracts using anti-PsaB antibody. Consistent with the predicted effects of cold temperature on PSI, PsaB amounts were significantly lower at 20°C than 27°C in all plant lines (**Figure 9A**). Pho1ΔL80 lines showed a more severe reduction in PsaB content compared to wildtype, BMF136, or the BMF136 transgenic lines expressing Pho1. Phenotypic characterization of the plants 150 to 180 DAT showed that Pho1ΔL80 plants were more sensitive to cold temperature (**Figure 9B**) as evident by increased yellowing of the leaves, burned leaf tips and a larger proportion of dead leaves. Analysis of the total dry biomass of the cold-threated plants also revealed reduced biomass for Pho1ΔL80 plants compared to wildtype and BMF136 (**Figure 9C**).

**Figure 9:**
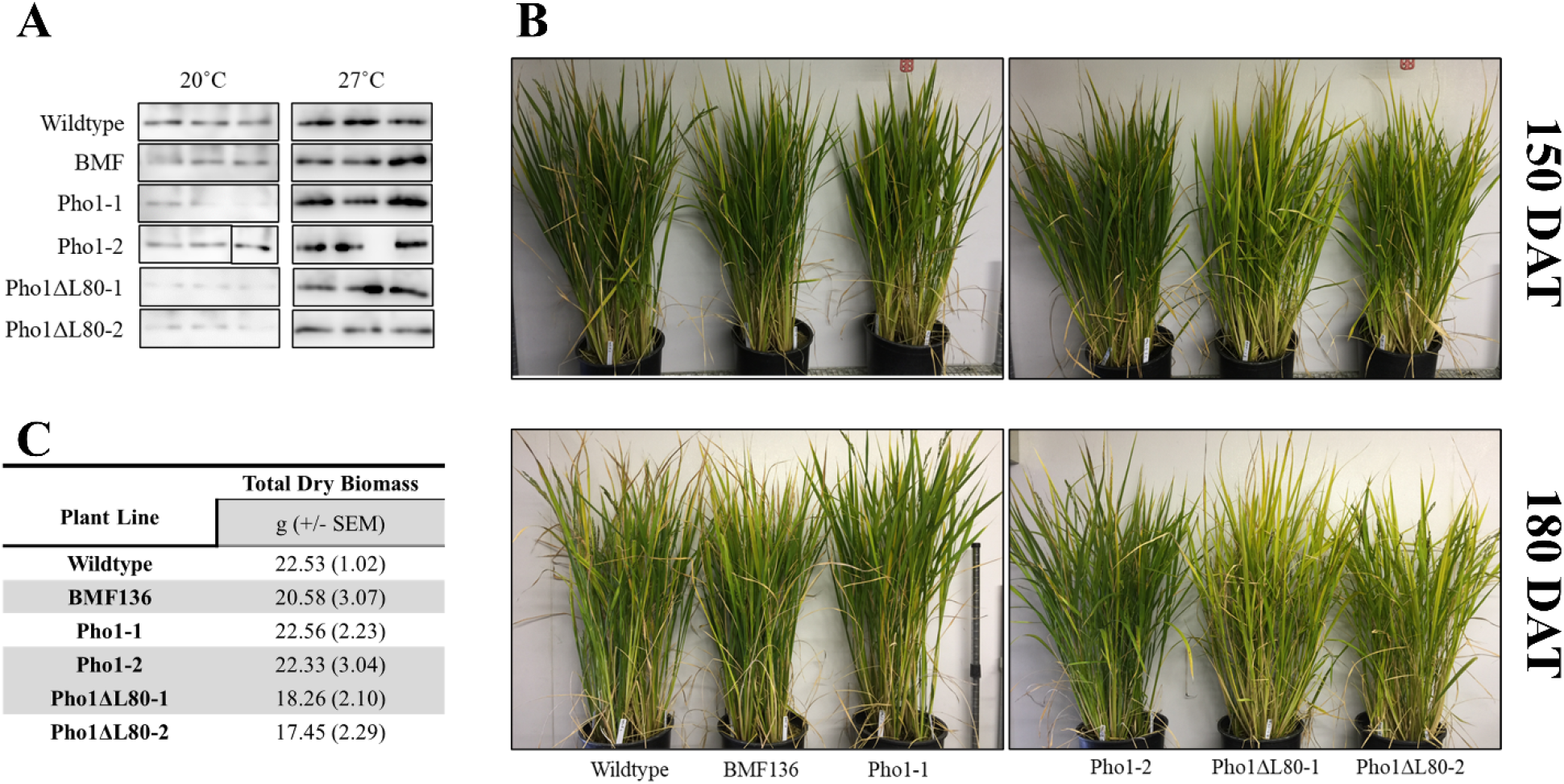
The response of wildtype, BMF136, Pho1, Pho1ΔL80 lines to cold stress. (A) The PsaB content of 120 DAT rice plants grown under normal (27°C) and cold (20°C) conditions. Leaf extracts (1 μg of chlorophyll) from each plant line was separated by SDS-PAGE and immunoblotted with anti-PsaB. (B) The growth phenotypes of plants 150 DAT and 180 DAT. (C) Biomass of wildtype, BMF mutant, and transgenic rice plants at the end of the trail under cold stress (210 DAT). Numbers of plants examined are wildtype (n=8), BMF136 (n=8), Pho1-1 (n=8), Pho1-2 (n=8), Pho1ΔL80-1 (n=8) and Pho1ΔL80-2 (n=8).

## Discussion

### Pho1 is a modulator of plant productivity and its L80 region acts as a regulator

Unlike the degradative role of glycogen phosphorylases, the plant Pho1 ortholog, has an established biosynthetic role in α-glucan (starch) metabolism [11, 13, 14, 17, 34–36]. In rice, the downregulation of Pho1 protein expression by mutations resulted in the production of grains ranging from shrunken to pseudo-normal in appearance but of smaller size. In this study, we complemented the *Pho1^−^* deficient rice mutant, BMF136, with Pho1 or Pho1ΔL80 to determine whether they could rescue the grain phenotype and to gain insight on the role of the L80 region. While complementation with Pho1 restored average grain weight to wildtype levels, expression of Pho1ΔL80 increased it above wildtype levels (**Table 1, Figure 2**), indicating a stimulation in starch synthesis and accumulation.

As removal of the L80 region does not affect the *in vitro* catalytic or regulatory properties of the rice Pho1 enzyme [21], the basis for the elevated starch accumulation (grain weight) remains to be resolved. Pho1 interacts with other starch biosynthetic enzymes including Dpe1 [21] and starch branching enzymes (SBE) [20]. The Pho1 enzyme complexes catalyze an alternative route of starch biosynthesis [1, 34, 37]. The major starch biosynthetic pathway involves the synthesis of ADPglucose from Glc1-P and ATP by ADP-glucose pyrophosphorylase and the subsequent transfer of the glucosyl moiety from ADP-glucose to an existing glucan primer by starch synthase activities [38, 39]. By contrast, starch synthesis by Pho1 is much simpler process as it can directly utilize Glc1-P to extend linear and branched α-glucans. This biosynthetic reaction by Pho1 is stimulated by branching enzymes as the latter activity effectively increases the substrate concentration by generating multiple branched non-reducing ends for the addition of glucosyl residues. Dpe1 stimulates Pho1 catalytic activity by elevating its affinity for substrate and by broadening its range for substrates [21]. The L80 peptide of the sweet potato Pho1 has been suggested to hinder substrate binding of α-glucans [40]. If the L80 peptide acts as a steric barrier in partially impeding the assembling of Pho1 with SBEs or Dpe1, the removal of the peptide from Pho1 can lead to more efficient formation of multi-protein complexes, which could result in increased biosynthetic activity.

A second possible causal basis for the elevated starch accumulation in developing grains of Pho1ΔL80 is that the source strength in rice leaves was increased to deliver more sugars to developing sink organs. In Arabidopsis and rice [22, 23, 41, 42], photosynthetic capacity and plant growth correlate with the levels of leaf starch synthesis [41, 42]. One underlying basis for this correlation is that starch synthesis serves as an effective alternative mechanism for recycling Pi needed for ATP synthesized by photophosphorylation. If Pho1ΔL80 stimulates starch synthesis in source green tissues, it would help maintain optimal ATP synthesis and increase the synthesis and levels of sugars that can be transported to sink organs for starch synthesis. However, like many other cereals, rice is naturally a poor leaf starch accumulator, hence its photosynthetic performance is not strongly influenced by transitory starch metabolism [38].

### Modulation of plant productivity by Pho1 can be an outcome of the stimulation of photosynthesis

The expression of Pho1ΔL80 also stimulates plant growth and development as evident by the increase in plant biomass and the earlier onset of panicle emergence and flowering. While the indirect influence of starch metabolism on photosynthesis could provide the basis for these enhanced phenotypic properties, the results presented in this study support a more likely explanation where Pho1 directly modulates photosynthesis. Pho1 interacts with PsaC as evident by the capture of the latter protein by anti-Pho1 and by the results obtained from the reciprocal pulldown down assays (**Figure 4 and 5**). Further evidence for this Pho1-PsaC interaction is that Pho1 is found associated with native PSI complexes of thylakoid membranes isolated from rice chloroplasts (**Figure 6**). Although Pho1 is considered a soluble stromal protein [14, 35, 36] (**Figure 5A**), its association with native PSI complexes infers that the enzyme’s *in vitro* interaction with PsaC, a peripheral subunit of the PSI complex located on the stroma side of the membrane [24], occurs *in situ*.

Analysis of the photosynthetic properties of wildtype, BMF136, Pho1, and Pho1ΔL80 transgenic lines indicates that the interaction between Pho1ΔL80 and PsaC has physiological relevance. Spectrophotometric studies of these plant lines *in vivo* readily demonstrates a significant change in the PSI properties of Pho1ΔL80 lines. Compared to wildtype and the other plant lines, Pho1ΔL80 plants showed a decrease in donor side limitation (YN_D_), and an increase in acceptor side limitation (YN_A_) **(Figure 7).** Two mutually exclusive hypotheses described below could explain the role of Pho1 and its L80 region on PSI.

### Hypothesis I: Pho1ΔL80 increases the efficiency of electron transport from PSI

Pho1ΔL80 plant lines show an increase in PSI acceptor side limitation YN_A_ (**Figure 7**) suggesting that the reduction of PSI acceptors is compromised in Pho1ΔL80 plant lines relative to wildtype, BMF136 and Pho1 lines. Such a higher YN_A_ implies that the flow of electrons from P700 (the primary donor of PSI) to PSI oxidized acceptors (PsaC, ferredoxin, NADP^+^, thioredoxin etc.) is impaired in Pho1ΔL80 **(Figure 10A)**. On the other hand, the higher acceptor site limitation may not be due to an inhibition of electron flow but is the result of a relative smaller pool of PSI acceptors in the oxidized state compared to that present in wildtype and the other plant lines **(Figure 10A)**. Assuming that the PSI acceptor pool sizes are not significantly different among the various plant lines, the smaller amounts of oxidized PSI electron acceptors in Pho1ΔL80 would correspond to a larger pool of acceptors in the reduced state. As the PSI properties were measured after adaptation of the plants to the various light intensities, the YN_A_ values represent a steady state condition and not the actual oxidation-reduction turnover rates of PSI acceptors. Hence, if the turnover rates (oxidized → reduced → oxidized) of PSI acceptors are comparable among the various plant lines, the larger reduced acceptor pool in Pho1ΔL80 would provide a greater source of reducing power and reductant to support (or drive) metabolism.

**Figure 10:**
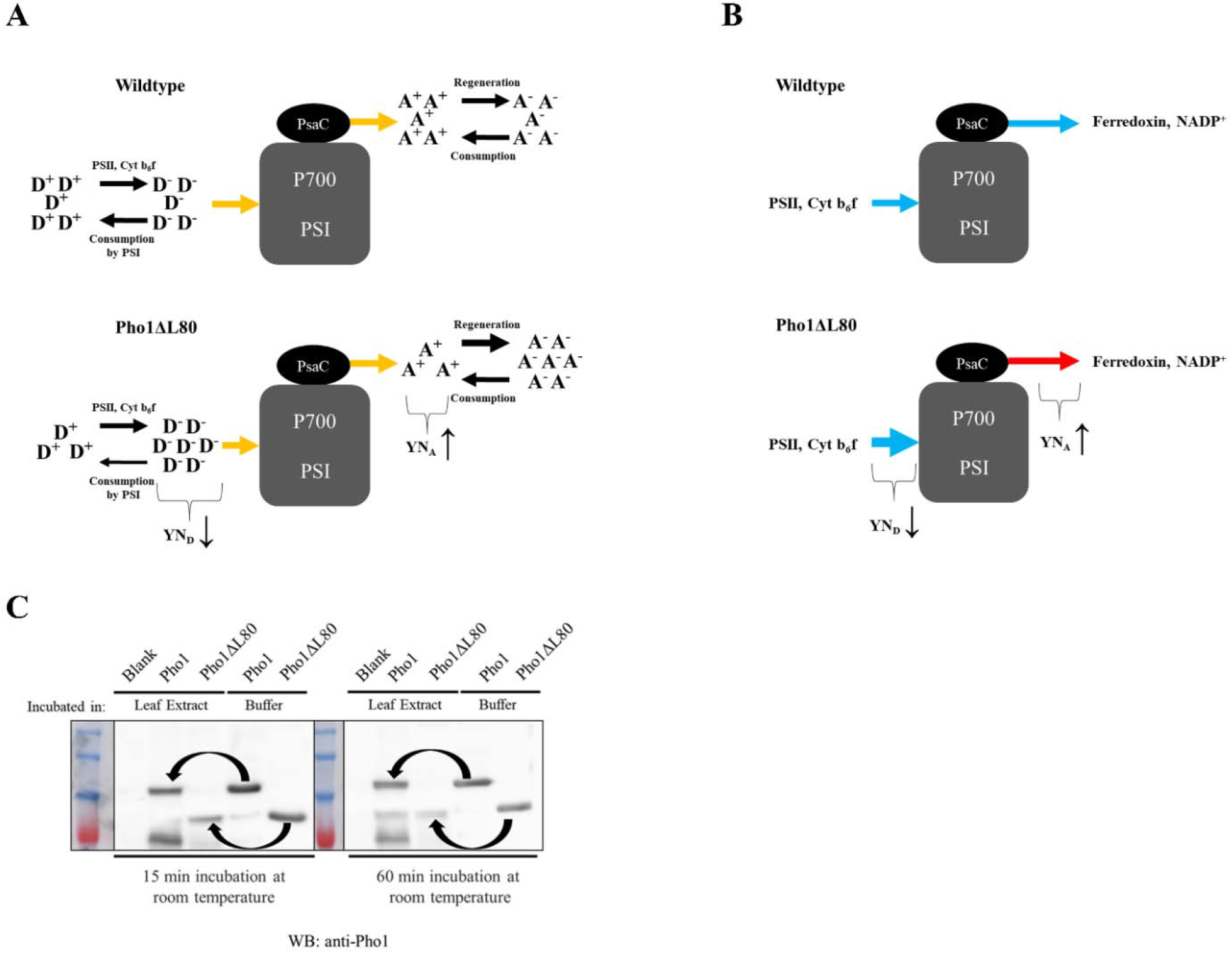
Changes in PSI properties of the Pho1ΔL80 plants compared to wildtype and a potential stability difference for the *in vivo* regulation of Pho1 and Pho1ΔL80. (A) Diagram explain PSI properties based on hypothesis I. Activities of PSII and cytochrome b6f regenerate the PSI donor pool, which is subsequently consumed by PSI. PSI, in turn, reduces oxidized acceptors and the reduced acceptors are used to drive metabolism. Availability of the reduced donor and oxidized acceptor pools determine PSI donor and acceptor site limitations. In Pho1ΔL80 plants, the acceptor pool is more reduced, but donor pool is more oxidized. This change is represented by lower YND and higher YNA in Pho1ΔL80 plants compared to wildtype. (B) Diagram explain PSI properties based on hypothesis II. In Pho1ΔL80 plants, electron flux to acceptors are more restricted (red arrow), while electron flow from donors is enhanced (thick blue arrow). This change is represented by lower YN_D_ and higher YN_A_ in Pho1ΔL80 plants compared to wildtype. (C) Stability of Pho1 and Pho1ΔL80 (2 μg each) after incubation with rice leaf extracts or with the extraction buffer for 15 min (left) or 60 min (right). Arrows point to the relative stability of Pho1 or Pho1ΔL80 incubated with leaf extracts compared to the extraction buffer. Blank is leaf extract without addition of recombinant Pho1 or Pho1ΔL80.

If “hypothesis I” is correct, the elevated reduced acceptor levels such as NADPH and reduced thioredoxin would impact other photosynthetic processes. Specifically, elevated reduced acceptor levels would more efficiently drive the C3 cycle and, in turn, CO_2_ assimilation. Such a condition would account for the higher CO_2_ assimilation rates exhibited by Pho1ΔL80 plants particularly at above ambient CO_2_ levels (**Figure 8**). Enhancing NADPH consumption was previously shown to increase photosynthetic efficiency and biomass production in cyanobacteria [43].

### Hypothesis II: Pho1ΔL80 inhibits electron flow from PSI, but increases the efficiency of electron transport to PSI

An alternative mechanism is that electron transport from PSI to its downstream acceptors is more restricted in Pho1ΔL80 plants without a change in the fraction of reduced/oxidized acceptors. Under this hypothesis, increased acceptor side limitation (**Figure 7**) observed for the Pho1ΔL80 plants is due to a reduction in intramolecular electron flow from P700 to ferredoxin and downstream acceptors NADP^+^, thioredoxin etc. However, since the PSI donor side limitation is also smaller and the quantum yield of photochemical energy conversion at PSI (YI) remained the same for Pho1ΔL80 plants, the inhibition of the acceptor side is compensated by the enhancement of the donor side (**Figures 7B, 7C and Figure 10B**).

If “hypothesis II” is correct, the enhancement of growth in Pho1ΔL80 plants could result from a higher rate of electron transport to PSI via the cyt b_6_f complex. In turn, ATP production is increased by the cyt b_6_f complex due to more proton pumping mediated by the higher electron transport rates upstream of PSI (lower YN_D_). Enhanced CO_2_ assimilation rates at higher CO_2_ concentration by Pho1ΔL80 plants **(Figure 8B)** could be a reflection of this property since CO_2_ assimilation under elevated CO_2_ makes the photosynthetic electron flux less sensitive to PSI acceptor side inhibition. This could indicate compensatory effects induced by Pho1ΔL80, for example, a change in cyt b_6_f complex abundance in thylakoid membranes.

### Pho1ΔL80 is more tolerant to nitrogen deficiency but more susceptible to cold stress

To determine a potential role for the L80 region, we examined the growth of Pho1ΔL80, wildtype, and other plant lines under nitrogen and phosphate deficiency. Interestingly, Pho1ΔL80 plant lines were much more tolerant to nitrogen deficiency than wildtype, BMF136 or the Pho1 transgenic lines. While overall growth of these various plant lines was lower under nitrogen deficiency compared to normal conditions, the Pho1ΔL80 lines grew 33-38% taller than wildtype under these conditions. By contrast, there was only an 18-23% difference in plant heights when Pho1ΔL80 and wildtype plants were cultured under normal nitrogen conditions. No significant differences in the changes in plant heights between Pho1ΔL80 and wildtype was evident between phosphate deficient and normal conditions.

Although Pho1ΔL80 outperformed wildtype under ideal conditions (**Figure 3**), they were more severely stressed under cold condition than the wildtype, BMF136, or Pho1 (**Figure 9**). Phenotypically, Pho1ΔL80 plants showed excessively yellowed leaves (**Figure 9B**) and tissue burning, which initiated at the tip of the leaf blade. This condition reduces the ability of the Pho1ΔL80 lines to maintain photosynthesis, accounting for their reduced total biomass at the time of harvest (**Figure 9C**). While all plant lines showed reduced amounts of PSI at 20°C (**Figure 9A**), consistent with the expected sensitivity of PSI to cold stress [32], the Pho1ΔL80 lines were the most affected.

Both hypothesis I and II can account on why the Pho1ΔL80 plants are more susceptible to cold. Hypothesis I assumes that under normal conditions the increased availability of reduced acceptors in Pho1ΔL80 plants drives metabolism and, in turn, plant growth. Under cold conditions, however, the demand for redox equivalents (such as Fd_red_ ^2-^, NADPH) is expected to decrease as metabolic processes that utilize this reducing power slow down due to lower temperature [44–46]. Such a condition would be especially severe for a naturally cold-sensitive plant like rice that has limited capacity to adjust its metabolism to cold. As recycling of reduced PSI acceptors is lowered, the availability of oxidized PSI acceptors further decreases. Reduction of PSI in the absence of available oxidized acceptors poses a serious problem because PSI has sufficient redox potential to reduce molecular oxygen and generate reactive oxygen species (ROS) when oxidized ferredoxin and NADP^+^ are not available [32, 47, 48]. If the Pho1ΔL80 plants have enhanced electron flow from PSI to downstream acceptors like the “hypothesis I’ describes, a decrease in the recycling of oxidized acceptors under cold could enhanced flow of electrons to oxygen generating ROS.

Alternatively under hypothesis II, the increased cold-sensitivity could be the result of the deceleration of the electron flux into PSI from the upstream components as the cyt b_6_f complex activity has a high temperature coefficient. Due to PSI’s capacity to generate ROS, plants regulate electron flow through PSI under conditions where the oxidized acceptor availability and recycling is low. This is achieved by two main mechanisms. First, linear electron flow to PSI is lowered to keep it oxidized as oxidized PSI can effectively dissipate light energy [48–50]. Second, cyclic electron flow is generated around the PSI to regenerate and maintain oxidized acceptors [48, 50]. If the decreased donor side limitation in the Pho1ΔL80 plants indicates an increased linear electron flow into PSI, the inability to control this under cold can become problematic. This is especially true if Pho1ΔL80 inhibits the electron flow from PSI to ferredoxin (as demonstrated by the higher YN_A_), while not influencing electron flow going into oxygen. Such a mechanisms would not only maintain a larger pool of reduced PSI due to enhanced linear electron flow, but it would also inhibit linear or cyclic flow out of PSI, leaving electron flow into oxygen as the only option and generating ROS.

In either case, resulting ROS and the damage to PSI is especially severe since PSI does not have an active repair mechanism and damaged PSI are usually degraded together with the associated pigments [32, 51]. Since the turnover rate of PSI subunits is relatively low, resulting ROS damage can cause prolonged photoinhibition of PSI activity [32]. In addition to PSI, ROS can cause widespread damage to other thylakoid membrane complexes [32, 47]. The buildup of ROS resulting from either one of these hypothetical mechanisms could be responsible for yellowing and burning of the rice leaf blades in cold-grown Pho1ΔL80 plants.

### The Role of Pho1 L80 region

The stimulation in photosynthetic performance exhibited by plants expressing Pho1ΔL80 (**Figure 7** and **Figure 8**) suggests that the L80 acts as a negative regulatory element in modulating PSI activity. While such regulation by L80 lowers photosynthesis and starch accumulation under normal growth settings, the peptide may be required under suboptimal growth conditions. Under low temperature growth conditions, the L80 region of Pho1 may be required in how Pho1 modulates PSI activity and diminishes the formation of ROS, and its deleterious effects on photosynthesis. Hence, the association of Pho1 with PsaC may play an active role in the photoinhibition mechanism of PSI when photosynthetic electron transport becomes imbalanced under suboptimal growth conditions [48].

### Potential regulatory elements in L80

Analysis of the L80 primary sequence identifies several potential regulatory motifs. One prominent one is the presence of a PEST motif located on the C-terminal half of the L80 region [1]. Although the functionality of the rice PEST motif is unknown, it can potentially alter the half-life of Pho1 compared to Pho1ΔL80. The sweet potato Pho1 was previously [52, 53] shown to be proteolyzed at its L80 region. Interestingly, incubation of recombinant Pho1 and Pho1ΔL80 protein in rice leaf extracts shows that Pho1ΔL80 degrades faster than Pho1 **(Figure 10C)**. This outcome is consistent with the previously reported role for L80 region of rice Pho1 in improving the heat stability of the enzyme [54]. It would be interesting to test if Pho1ΔL80 is also degraded faster with chloroplast extract that lacks proteasomes, but technical difficulty of collecting large quantities of intact chloroplast from rice tissue makes this study difficult.

Several residues of the maize [55] and sweet potato [40] L80-like domain are phosphorylated. Although phosphorylation of the rice Pho1 L80 has not been demonstrated, the phosphorylated Ser residues of the maize Pho1 [55] are conserved in the rice L80. Such post-translationally modified Ser on Pho1 L80 can be part of a binding motif for 14-3-3 proteins, which are known to interact with phosphorylated Ser and Thr residues [56, 57]. Binding of the 14-3-3 to their target proteins can alter the conformation, activity, stability, or intercellular localization [58, 59]. 14-3-3 proteins are known to influence grain filling in rice [60, 61], and several starch biosynthetic enzymes such as starch synthase are known targets of 14-3-3 proteins [62, 63]. It has been proposed that the dimeric 14-3-3 can act as scaffolding proteins by binding to multiple phosphorylated starch biosynthetic enzymes and keeping them together in a phosphorylation dependent manner [64]. Similarly, the association of 14-3-3 with phosphorylated L80 may attract other proteins that regulate PSI activity.

Computational analysis [57] suggest certain serine and threonine residues upstream and downstream of L80 could be targeted by 14-3-3 protein **(Supplementary Figure 3)**. Interestingly, this analysis does not predict any residue on L80 region as a target. Nevertheless, L80 region could have a role in a potential 14-3-3 regulation through conformational changes or through influencing interacting partners.

## Conclusion

Enhanced photosynthetic performance and growth of Pho1ΔL80 plants suggests that Pho1 has a role in modulating photosynthesis with L80 acting as a regulator. The cold sensitivity of Pho1ΔL80 plants supports the view that the L80 is essential under conditions where metabolic flexibility is diminished. As the L80 region of Pho1 does not influence the catalytic properties of the enzyme, it is unlikely that the observed phenotype is related to Pho1’s enzymatic role. Nevertheless, the L80 suppresses Pho1’s role in starch biosynthesis. Although L80 may simply acts as a steric barrier in reducing the ability of Pho1 to associate with other starch biosynthetic enzymes, there may be other starch-related processes that require the L80 of Pho1.

While substantial evidence indicate that Pho1 interacts with PsaC and may modulate PSI activity, the molecular mechanism of how Pho1 and its L80 region influence PSI activity is unclear. Two mutually exclusive hypotheses **(Figure 10A and 10B)** can account for the changes in the PSI properties exhibited by Pho1ΔL80 plants. Both hypotheses suggest Pho1 influence the electron flow through PSI, but they disagree on the underlying basis. According to hypothesis I, in the absence of the negative L80 modulator, Pho1ΔL80 possesses a larger pool of reduced PSI acceptors that can drive C3 metabolism. On the other hand, hypothesis II suggests that Pho1ΔL80 simultaneously inhibit the flow of electrons out of PSI to acceptors while enhancing electron flow into PSI. *In vitro* studies that specifically investigate PSI activity in the absence or presence of Pho1/Pho1ΔL80 could shed light on the underlying mechanisms and provide support for one of the two hypotheses.

One curious observation is that although BMF136 (*pho1^−^* mutant) line matures slower than wildtype, its photosynthetic properties are not significantly different from wildtype. BMF136 is a Pho1 knockdown mutant that produces a variant protein that has a small deletion of 56 residues located in the N-terminal half of the primary sequence as well as apparently very small amounts of wildtype Pho1 subunit **(Figure 1B)**. As the native Pho1 exists as a dimer, the two Pho1 subunit types can potentially form three different dimeric forms, two of which are likely non-catalytic. The smaller Pho1 variant polypeptide retains much of the normal primary sequences including the L80 and may still be functional in interacting with PsaC and, in turn, modulating PSI activity. Hence, it would be interesting to see whether a rice line harboring a *pho1^−^* “knockout” mutation exhibits lower photosynthetic properties than wildtype.

## Materials and methods

### Plasmid construction and rice transformation

Rice transformation vectors pSH694 and pSH695 were generated by cloning the RuBisCO small subunit chloroplast transit peptide sequences in frame to the mature rice Pho1 or Pho1ΔL80 cDNA sequence [54] under the control of a native Pho1 promoter into pCambia1300. Rice transformation vectors pKK29 and pKK30 were constructed by replacing the Pho1 promoters and RuBisCO sequences with rice AGPase small subunit 2 (AGPS2) promoter and the first exon and intron of rice Pho1 harboring native chloroplast transit peptide, respectively. Backbone vectors for pKK29 and pKK30 constructions were pCambia1300 and pCambia1301, respectively. The AGPS2 promoter was selected due to its strong expression profile and similar tissue specificity compared to native Pho1 promoter [12]. These vectors were then transformed into the Pho1-deficient (*Pho1^−^*) mutant rice BMF136 (derivative of *Oryza sativa* L.cv TC65) [11] using *Agrobacterium tumefaciens* AGL1 [65] and screened with hygromycin. Homozygous plants expressing Pho1 or Pho1ΔL80 were identified by western blot analysis using rice Pho1 antibody [21]. Homozygous lines (T3 generation) of each transgenic line, the parental line BMF136, and wildtype TC65 were used in this study.

### Plant growth conditions

Rice plants used in this study were grown in parallel, in environment-controlled chambers with ambient CO_2_. The photoperiod was set to 12 h of day and 12 h of night with a relative humidity of 70% and a day temperature of 27°C and a night temperature of 23°C. Light intensity was set to 650 μmol m^−2^ s^−1^ at the midpoint between the canopy and the base of the plant. Plants were fertilized twice a week throughout the growth period.

Plants subject to cold stress were grown under the same conditions described above except a constant day and night temperature of 20°C.

Plants subjected to phosphorus or nitrogen deficiency were grown under the same conditions described above, except being watered with premixed fertilizer solutions prepared in distilled water lacking the respective nutrient [66]. Control plants were watered daily with complete fertilizer solution [10 mM Ca(NO_3_)_2_, 10 mM KNO_3_, 4 mM MgSO_4_, 2 mM KH_2_PO_4_] plus micronutrient solution (93 μM H_3_BO_3_, 18 μM MnCl_2_, 1.6 μM ZnSO_4_, 0.64 μM CuSO_4_, 0.25 μM Na_2_MoO_4_) and 6 mg/L Sequestrene 330 Fe (Sprint^®^). Plants investigated for phosphorus deficiency were watered daily with a premixed fertilizer solution lacking phosphate, and plants investigated for nitrogen deficiency were watered daily with a premixed fertilizer solution lacking nitrate.

### Phenotypic analysis of plants

Mature grains were harvested from the plants and air-dried for 7 days at room temperature. Individual grain weights were determined and averaged for calculating average grain weight for each line. Individual grain weights were also used for determining grain weight distribution. The total grain weight per plant was determined by weighing all grains from each plant. The number of panicles for each line was counted and averaged to calculate the average panicle number. The number of grains per panicle was calculated by dividing the total grain weight by average grain weight, and then dividing the resulting value by average number of panicles.

### Protein extraction from plant tissues

A slightly different protein extraction procedure was used depending on the investigated rice tissue. For mature rice grains, dehusked grains were first powdered and proteins were extracted by incubating the grain powder in sufficient amount urea/SDS extraction buffer (50 mM Tris-HCl, pH 7.0, 6.4 M urea, 2% SDS, 1% β-mercaptoethanol, 5% glycerol and 0.02% bromophenol blue) on ice for 10 min. Softer tissues such as developing grains, green seedlings (minus roots), leaves, and sheaths were first sliced into a manageable size, crushed in enough urea/SDS buffer using a pestle and incubated on ice for 10 min. Next, samples were centrifuged at 12,000 RPM and the supernatant fractions containing the extracted proteins were transferred to new tubes.

### SDS-PAGE

SDS-PAGE analysis of proteins was performed as described [67]. Briefly, samples containing native Pho1, transgenic Pho1 or Pho1ΔL80, and recombinant Pho1 or Pho1ΔL80 proteins were separated on a 7.5% SDS-polyacrylamide gel with Tris-glycine running buffer (25 mM Tris, 192 mM glycine, 0.1% SDS, pH 8.3). PsaB proteins were separated on a 10% SDS-polyacrylamide gel with the same buffer, and PsaC proteins were separated on a 12% SDS-polyacrylamide gel with Tris-Tricine running buffer (anode buffer: 100 mM Tris-HCl, pH 8.9; cathode buffer: 100 mM Tris, 100 mM Tricine 0.1% SDS, pH 8.3).

### Immunoblot analysis

Proteins separated on SDS polyacrylamide gels were transferred onto 0.2 μm nitrocellulose membrane (Pall Corporation). The blotted membrane was blocked with 5% (w/v) non-fat milk in 1X TBS (50 mM Tris-HCl, pH 7.4, 150 mM NaCl) and then probed in the same solution at 4°C with the following primary antibodies: anti-Pho1 (1/2000 dilution) [21], anti-PsaC (PhytoLab PHY0055A, 1/500 dilution), anti-PsaB (PhytoLab PHY0054A, 1/1500 dilution). Afterwards, the membrane was washed 4 times with 1X TBS containing 0.1% (v/v) Tween-20 and incubated with 1/5,000 diluted goat anti-rabbit horseradish peroxidase-conjugated secondary antibody (Thermo Scientific) in 5% milk TBS for 1 hour at room temperature with shaking. The membrane was then washed 4 times with 1X TBS with 0.1% (v/v) Tween-20. The antibody-antigen complex was detected by using the SuperSignal West Pico PLUS Chemiluminescent Substrate (Thermo Scientific) and the FujiFilm LAS-3000 or Amersham imager 600 analyzers.

### Silver staining

Silver staining of the gel was performed as described [68] with minor modifications. Briefly, the SDS-PAGE gel was fixed in in solution containing 50% methanol and 5% acetic acid for 20 min and washed with 50% methanol and water for 10 min each. The gel was then incubated in sensitizing solution (0.02% w/v sodium thiosulfate in water) for 1 min and washed two times with water for 5 min each. The staining reaction was initiated by incubating the gel in silver solution (0.1% w/v silver nitrate, 0.08% v/v formaldehyde) and then washing twice with water for 1 min. The gel was then incubated in a solution containing 2% w/v sodium carbonate and 0.04% formaldehyde and then in 5% acetic acid.

### Co-immunoprecipitation (Co-IP)

Polyclonal rabbit antibodies generated against Pho1 [21] were affinity-purified as described [69]. All steps were performed at 4°C unless otherwise indicated. Twenty μL of protein A-agarose slurry (Invitrogen) was centrifuged at 1,000 X g for 1 min and washed two times with PBS [137 mM NaCl, 2.7 mM KCl, 10mM Na_2_HPO_4_, 1.8 mM KH_2_PO_4_]. Twenty μg of affinity-purified anti-Pho1 antibody (or IgG, control) in PBS was allowed to bind overnight to Protein A agarose beads. The agarose beads were collected by centrifugation at 1,000 X g for 1 min and then washed three times with 500 μL PBS to remove unbound antibodies. Protein A–antibody complex was crosslinked by incubating the beads in 100 μL PBS containing 0.45 mM disuccinimidyl suberate (DSS) at room temperature for 1 h with constant mixing. At the end of the crosslinking, the solution was removed by centrifugation at 1,000 X g for 1 min and free antibodies were removed by washing the beads two times with 50 μL of 100 mM glycine buffer, pH 2.5, and two times with PBS. Protein A-agarose beads with crosslinked anti-Pho1 (or IgG) antibody were equilibrated with protein extraction buffer [20 mM Tris-HCl pH 7.5, 150 mM NaCl, 1 mM EDTA, 2 mM ATP, 2 mM NaF, 1 mM dithiothreitol (DTT), 1 mM phenylmethylsulfonyl fluoride (PMSF), 1X protein inhibitor cocktail and 0.5% Nonidet P-40]. In parallel, 3 g of plant tissue (developing grains or grainlings) were ground in 3 ml of the same protein extraction buffer. Tissue extracts were centrifuged at 1,500 X g and then at 15,000 X g to remove starch and cell debris. One-ml of soluble tissue extract was applied to Protein A-agarose beads with crosslinked anti-Pho1 antibody (or IgG) and incubated 2-4 h at 4°C with constant mixing. The mixture was then centrifuged 1,000 X g for 1 min to collect the beads, which were subsequently washed twice with 500 μL of wash buffer (protein extraction buffer minus Nonidet P-40) and once with 500 μL PBS to remove unbound proteins. Proteins initially captured by the anti-Pho1 beads were eluted with 20 μL of the low pH glycine buffer for 5 min at room temperature followed by a second elution with 60 μL of the low pH glycine buffer for 10 min at room temperature. Eluted samples were immediately neutralized with 1 M Tris-HCl, pH 8.8. Proteins co-immunoprecipitated with anti-Pho1 or by control IgG were analyzed by liquid chromatography-tandem mass spectrometry (Washington State University Tissue Imaging and Proteomics Laboratory). Proteins interacting with Pho1 were determined by differentiating identified proteins in the anti-Pho1 eluate against those in the IgG control eluate.

### Protein purification

OsPho1, OsPho1ΔL80, HALO-Pho1, HALO-Pho1ΔL80, HALO-PsaC and HALO-GFP were expressed and purified as described [18, 21, 54, 70]. Briefly, protein expression constructs were grown in *Escherichia coli* EA3457 strain and. protein expression was induced overnight with 0.15 mM of IPTG when the OD_600_ was between 0.8 to 1.0 at room temperature for OsPho1, OsPho1ΔL80, and at 16°C for HALO-Pho1, HALO-Pho1ΔL80, HALO-PsaC and HALO-GFP. The bacterial cells were collected by centrifugation and lysed by sonication. After clarification of the extract by centrifugation at 12,000 RPM, the proteins were purified to near homogeneity by DEAE-Sepharose FF (GE Healthcare) chromatography followed by TALON Superflow (GE Healthcare, a cobalt-based IMAC resin) affinity chromatography.

### Halo-protein pulldown

Halo-linked PsaC (HALO-PsaC, pKK32) was generated by PCR amplification of the PsaC gene from rice chloroplast DNA and by cloning in-frame next to the Halo-tag sequences on pSH583 [54]. Halo-linked Pho1 (Halo-Pho1, pSH585 [54]), Halo-linked Pho1ΔL80 (Halo-Pho1ΔL80, pSH642 [54]), Halo-linked GFP (Halo-GFP, pSH583 [54]) and Halo-PsaC were expressed and purified as described above. Purified Halo-proteins were used for HaloTag^®^ pulldown assays as described [54]. 100 μL of HaloLink slurry (Promega) was centrifuged at 800 X g for 1 min to collect the resin. The supernatant was discarded, and the resin was washed three times with 800 μL of binding buffer (25mM HEPES-NaOH pH 7.5, 150mM NaCl, 0.05% Nonidet P-40). The equilibrated resin was collected by centrifugation at 800 X g for 2 min. 500 μg of the purified HALO-protein was added to the HaloLink resin and incubated at 4°C for 2 h to allow binding. Unbound proteins were removed by centrifugation at 800 X g for 2 min and the resin washed two times with binding buffer and two times with the protein extraction buffer (see above). In parallel, 3 g of whole green seedling tissue were ground in 3 ml of the protein extraction buffer. Tissue extracts were centrifuged at 1,500 X g and then at 15,000 X g to remove starch and cell debris. One ml of soluble tissue extract was applied to the HaloLink resin coupled to Halo-Pho1, Halo-Pho1ΔL80, Halo-PsaC or Halo-GFP and incubated for 2 h at 4°C, followed by 15 min incubation at room temperature. The mixture was centrifuged at 800 X g for 2 min to collect the resin, which was subsequently washed three times with wash buffer (20 mM Tris-HCl pH 7.5, 150 mM NaCl, 2 mM NaF, 1 mM DTT). Bound proteins were eluted from the resin by incubating with 100 μL of Urea/SDS extraction buffer (50 mM Tris-Cl pH 7.0, 6.4 M urea, 2% SDS, 1% β-mercaptoethanol, 5% glycerol and 0.02% bromophenol blue) at room temperature for 10 min. Proteins captured by the Halo-tagged bound proteins were collected by centrifugation at 800 X g for 2 min.

### Isolation of rice chloroplasts

Chloroplasts were isolated from rice seedlings as described [27] with the following modifications. All steps were performed at 4°C unless otherwise indicated. Twenty g of green seedling tissue was powdered in liquid nitrogen and homogenized in cold isolation buffer (50 mM HEPES pH 7.8, 330 mM sorbitol 5 mM MgCl_2_, 2 mM EDTA, 2 mM NaF). Homogenate was filtered through four layers of Miracloth (Millipore). The filtrate was then centrifuged at 200 X g for 3 min to remove cell debris. The supernatant fluid was then transferred to a new tube and centrifuged at 1,000 x g for 7 min to sediment crude chloroplasts. Chloroplast pellet was resuspended in 6 ml of the isolation buffer containing 0.1% w/v BSA. Ten ml of 40% Percoll (Pharmacia) in the isolation buffer was prepared. 6 ml of the chloroplast suspension was layered on top of the 10 ml 40% Percoll in the isolation buffer and centrifuged at 1,700 X g for 7 min. The intact chloroplasts formed a small green pellet at the bottom of the tube. After the Percoll solution containing broken chloroplasts was carefully removed, the intact chloroplast pellet was washed twice with the isolation buffer and solubilized in 500 μL of the isolation buffer. Chlorophyll concentration of the isolated chloroplasts was determined in 80% acetone by using the Porra method [71].

### Isolation of rice thylakoid membrane complexes

Thylakoid membrane complexes were isolated as previously described [27]. All steps were performed at 4°C unless otherwise indicated. Chloroplast suspension containing 100 μg of chlorophyll was lysed by incubating in hypo-osmotic buffer (50 mM HEPES pH 7.5, 2 mM MgCl_2_, 1 mM EDTA, 1 mM NaF, 1mM PMSF) for 30 min at 4°C. Thylakoid membranes were collected from the lysate by centrifugation at 15,000 X g for 15 min. Sedimented thylakoid membranes were solubilized in 100 μL of ACA/DDM buffer (750mM aminocaproic acid, 50 mM Bis-Tris, pH 7.0, 0.5 mM EDTA, 4% n-Dodecyl β-D-maltoside). The homogenate was centrifuged at 15,000 X g for 30 min and the supernatant was transferred to a new tube containing 20 μL of CBB buffer (5% Coomassie brilliant blue G250 in 750 mM aminocaproic acid). The samples were then separated on blue native gels to visualize thylakoid membrane complexes.

### Blue native PAGE

Blue native PAGE was conducted as previously described [72]. Bis-Tris acrylamide gels (Invitrogen NativePAGE™) consisting of a continuous 4-16% acrylamide gradient were used for the separation. Fifteen μL of solubilized thylakoid membrane (equivalent to 80 μg chlorophyll) was loaded onto each lane. Gel filtration protein standards (Pharmacia) of thyroglobulin (669 kDa), ferritin (440 kDa) and catalase (232 kDa) were used as native molecular weight markers. Gels were run for ~20 h at 4°C using an anode buffer of 50 mM Bis-Tris (pH 7.0) and a cathode buffer of 50 mM tricine, 15 mM Bis-Tris (pH 7.0), 0.02% CBB-G250. Separated protein complexes could be directly visualized without staining.

### Extraction of thylakoid membrane complexes from blue native gels

All steps were performed at 4°C unless otherwise indicated. Bands of interest containing the thylakoid membrane complexes were excised from the blue native gels and transferred to capped tubes. 7-10 stainless steel beads (1.8-2 mm in diameter) were added to each tube. Samples were frozen at −80°C for 20 min and then transferred to liquid nitrogen. Frozen samples were pulverized by a bead homogenizer (Omni Bead Ruptor 24 Elite) for 30 sec at a setting of 5 ms. One-half ml of ice-cold 1X SDS sample buffer (50 mM Tris-Cl pH 7.0, 2% SDS, 1% β-mercaptoethanol, 5% glycerol and 0.02% bromophenol blue) was added to each tube. Samples were vibrated for an additional 30 sec at 5 ms and the tubes were incubated on ice for 20 min to allow protein diffusion. Samples were centrifuged at 5,000 RPM for 10 min and the supernatant transferred to a new tube. To collect the remaining solution inside the gel slurry, the metal beads were removed and the gel slurry was passed through a filtration column (Bio-Rad Econo-Pac column) by centrifugation at 8,000 RPM for 10 min. Flow-through fractions from the column were combined with the previously collected supernatant and the 1X SDS sample buffer was used to adjust the samples to equal volumes.

### Dot-blot analysis

10 μL each of the thylakoid membrane proteins extracted from blue native gel bands was spotted on a nitrocellulose membrane (Pall Corporation). After spots on the membrane were completely air-dried, Pho1 protein was detected using affinity-purified anti-Pho1 as described above.

### Stability assay of Pho1 and Pho1ΔL80 in leaf extract

Four g of rice leaves were homogenized in 100 ml extraction buffer (20 mM Tris-Cl pH 7.5, 150 mM NaCl, 1 mM DTT, 1 mM ATP, 0.5% Nonidet P-40), and filtered through four layers of Miracloth (Millipore). The filtrate was centrifuged at 200 X g for 3 min to remove cell debris, and the soluble leaf extract was then transferred to a new tube. In order to assess the degradation of Pho1 and Pho1ΔL80 in the leaf extract, two μg of the purified recombinant Pho1 or Pho1ΔL80 was added to a microcentrifuge tube. One-hundred μL of leaf extract was added on top of the recombinant proteins and incubated for 15 or 60 min at room temperature with gentle shaking. In parallel, the extraction buffer was added instead and incubated under the same conditions At the end of the incubation, the SDS sample buffer was added to each mixture and the proteins are immediately separated by SDS-PAGE, transferred to nitrocellulose membrane and immunoblotted with anti-Pho1. The degradation level of Pho1 or Pho1ΔL80 was assessed by comparing the intensity of the Pho1 or Pho1ΔL80 bands from the leaf extract treatment with that of the extraction buffer-treated Pho1 or Pho1ΔL80.

### Determination of photosystem-I functionality

P700 redox states is measured with intact rice plants from 820 nm minus 900 nm absorption changes using a home-built flash-spectrometer with ms time-resolution [73]. From the P700 redox changes the quantum yield of photochemical energy conversion at PSI (YI) and quantum yields of non-photochemical energy dissipation due to donor side (YN_D_) or acceptor side (YN_A_) limitations, were derived at different light intensities as described [30]. Rice plants were used 28-35 days after transplantation (DAT) for the study. Measurements were conducted on the widest section of the youngest fully exposed leaf on six to eight different plants (n = 12 to 32) after dark-adaptation at room temperature.

### Gas exchange measurements

Gas exchange measurements were conducted to calculate photosynthetic parameters as previously described [74] using LI-6400XT portable photosynthesis system (LI-COR Biosciences). Rice plants used were 28-49 DAT. Measurements were conducted at the widest section of the youngest fully exposed leaves of six to fourteen different plants (n = 6 to 14) after dark acclimation. Two leaf blades from each plant were used to cover the cuvette surface. Measurements were conducted after plants were acclimated and photosynthetic parameters stabilized. The data was collected between 9 _AM_ and 5 _PM_ in a chamber with an air temperature around 27°C.

## Acknowledgments

We thank Dr. Rita Giuliani for technical assistance.

**Supplementary Data.** Pho1 captured proteins identified by LC-MS/MS

---

**Supplementary Figure 1.**
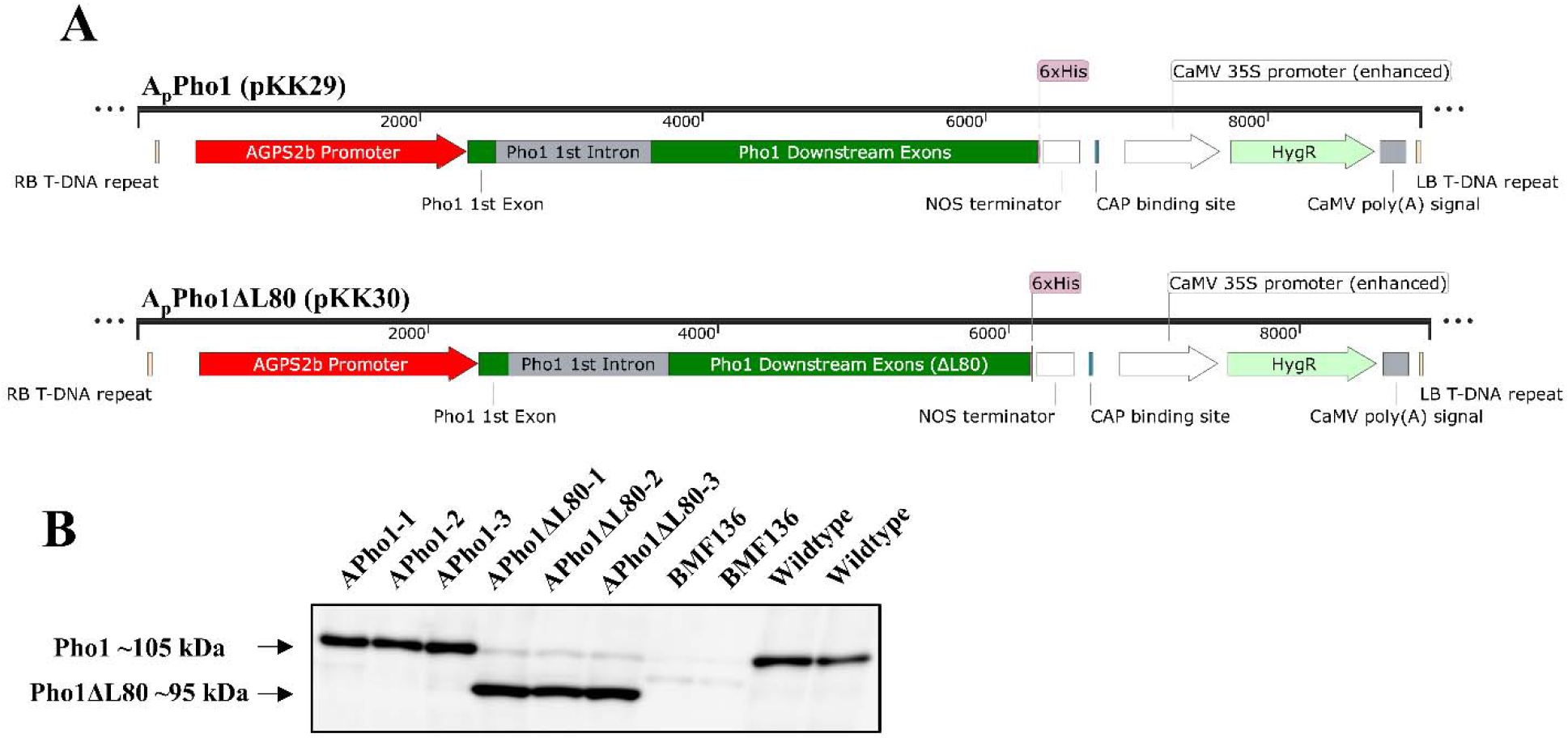
AGPS2 promoter-driven expression of Pho1 and Pho1ΔL80 in Pho1-deficient BMF136. (A) Maps (created by SnapGene) of DNA constructs introduced into BMF136. (B) Representative expression levels of Pho1 or Pho1ΔL80 in mature seeds of transgenic rice lines. One seed from each plant was crushed and proteins extracted using 200μL of urea/SDS extraction buffer. 7.5 μL of the sample was loaded on to a 7.5% SDS polyacrylamide gel, separated and transferred to a nitrocellulose membrane. Anti-Pho1 antibody was used to examine the relative expression levels of Pho1 (105 kDa) and Pho1ΔL80 (95 kDa).

**Supplementary Figure 2.**
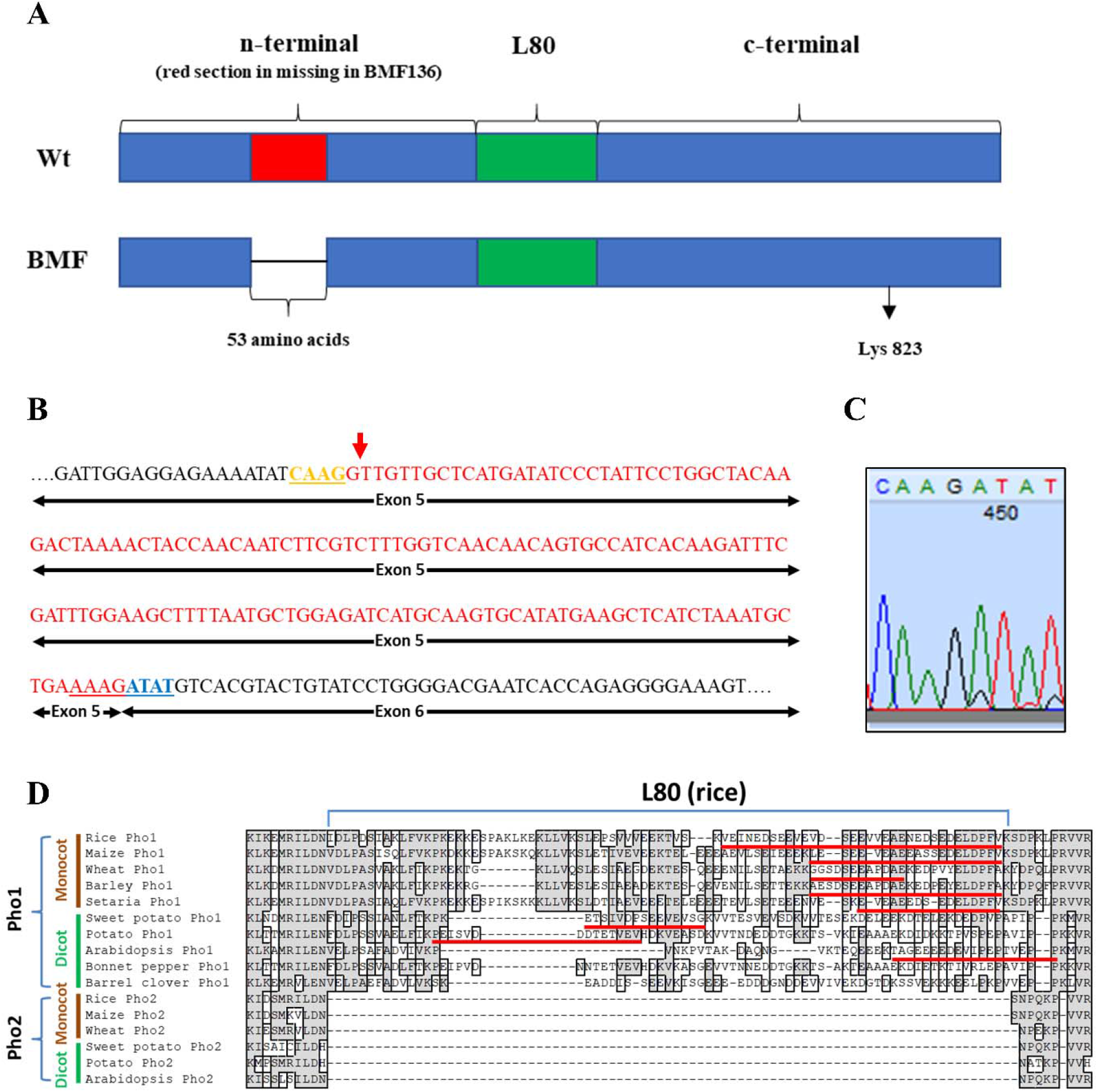
Pho1 transcript of BMF136 and the alignment of L80 regions of rice and various other plant species. **A.** Diagram showing deletion in BMF136 due to improper splicing. Red region shows the spliced-out section of the 5^th^ exon in BMF136, green is the L80 region, Lys823 is the Schiff-base lysine. Deletion is in frame and therefore does not cause a frameshift. **B.** Red shows the spliced-out section of the 5^th^ exon in BMF136 cDNA. Marked GT site at the 5’ of the red sequence is improperly used as the splicing site, as the mutation in BMF136 changed the true GT site on the downstream intron. Sequences shown in yellow and blue are the sites spliced together in BMF136. **C.** DNA chromatogram of Pho1 cDNA sequence from BMF136 showing the joining of yellow and blue sequences from panel B. **D.** L80 regions of Pho1 from various plants aligned with respect to each other, and the Pho2 sequence. Red lines show the predicted PEST motif.

**Supplementary Figure 3.**
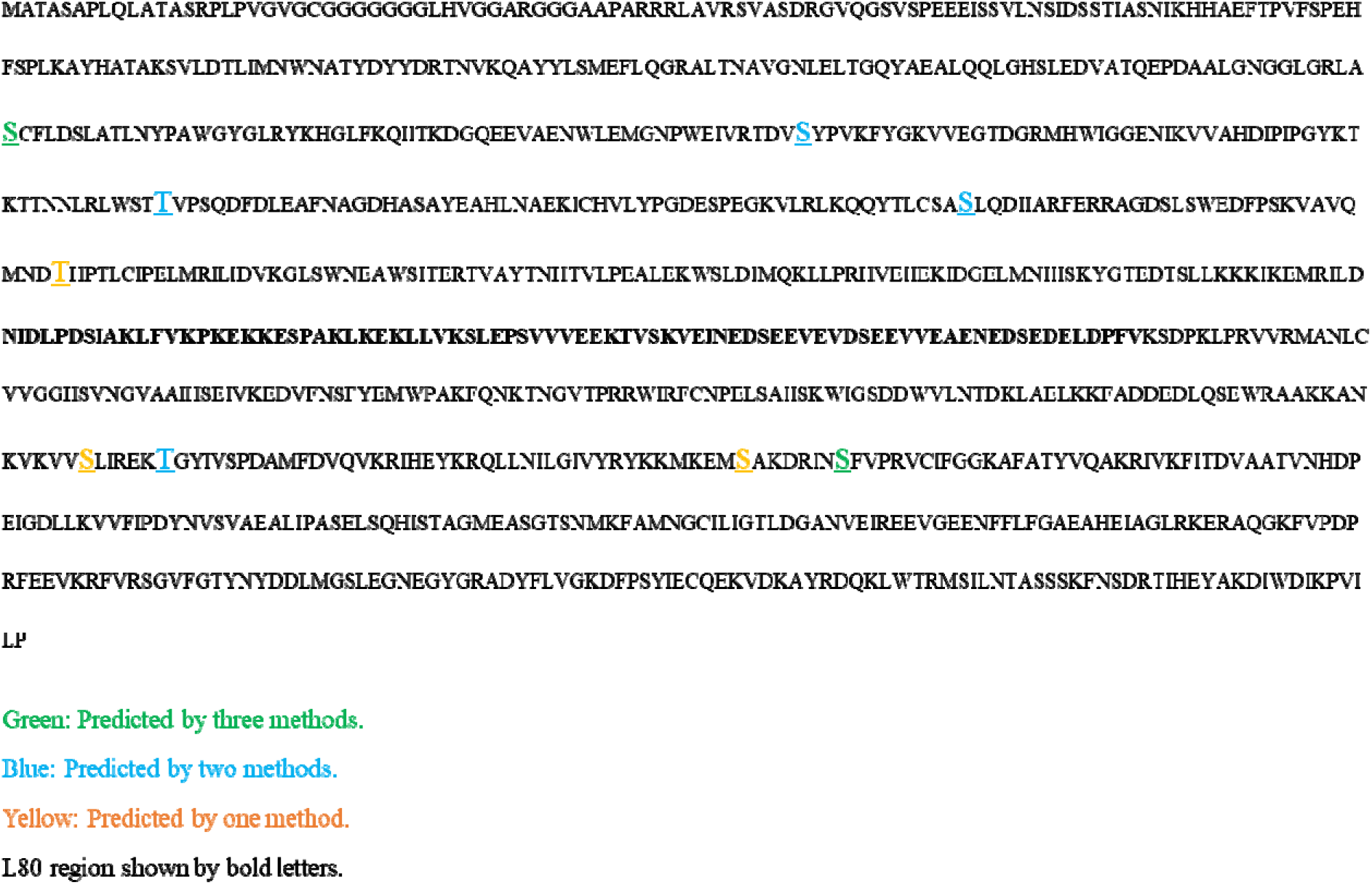
Predicted 14-3-3 targeted sequences on rice Pho1. Sequences targeted by 14-3-3 proteins are computationally predicted by the 14-3-3-Pred server. Green show residues predicted by 3 methods, blue shows residues predicted by two methods and yellow show residues. L80 region is shown with bold letters.

**Supplementary Data.** Pho1 captured proteins identified by LC-MS/MS

